# Chromatin remodeling protein BPTF regulates transcriptional stability in planarian stem cells

**DOI:** 10.1101/2024.05.24.595819

**Authors:** Prince Verma, Alejandro Sánchez Alvarado, Elizabeth M. Duncan

**Affiliations:** Department of Biology, University of Kentucky, Lexington KY, USA; Stowers Institute for Medical Research, Kansas City MO, USA

## Abstract

Trimethylation of histone H3 lysine 4 (H3K4me3) correlates strongly with gene expression in many different organisms, yet the question of whether it plays a causal role in transcriptional activity remains unresolved. Although H3K4me3 does not directly affect chromatin accessibility, it can indirectly affect genome accessibility by recruiting the ATP-dependent chromatin remodeling complex NuRF (Nucleosome Remodeling Factor). The largest subunit of NuRF, BPTF/NURF301, binds H3K4me3 specifically and recruits the NuRF complex to loci marked by this modification. Studies have shown that the strength and duration of BPTF binding likely also depends on additional chromatin features at these loci, such as lysine acetylation and variant histone proteins. However, the exact details of this recruitment mechanism vary between studies and have largely been tested in vitro. Here, we use stem cells isolated directly from live planarian animals to investigate the role of BPTF in regulating chromatin accessibility in vivo. We find that BPTF operates at gene promoters and is most effective at facilitating transcription at genes marked by Set1-dependent H3K4me3 peaks, which are significantly broader than those added by the lysine methyltransferase MLL1/2. Moreover, BPTF is essential for planarian stem cell biology and its loss of function phenotype mimics that of Set1 knockdown. Together, these data suggest that BPTF and H3K4me3 are important mediators of both transcription and in vivo stem cell function.

## INTRODUCTION

Planarian flatworms are a robust and powerful model for studying in vivo stem cell function. Not only do planarians maintain adult stem cells throughout their lifetime, but each worm also exhibits the fundamental properties of a stem cell: they can both regenerate large areas of complex tissue and, at the same time, renew their regenerative capacity so that the same animal can repeat this incredible feat again [1]. Planarian regeneration is fueled by a large population of multi and pluripotent stem cells (also known as neoblasts) that proliferate and differentiate in response to injury signals [2]. In addition to their essential role in tissue regeneration, planarian stem cells have many other remarkable characteristics. For example, they have a strikingly high tolerance for ionizing radiation, proliferate indefinitely without showing signs of senescence or exhaustion and, despite these other features, rarely form tumor-like structures [3–6]. The planarian stem cell population is also transcriptionally heterogeneous [7–13], with subsets of stem cells expressing various lineage-specific genes in addition to established planarian stem cell markers (e.g., *piwi-1*). Several studies have shown that these lineage-marked stem cells are required for the regeneration of their associated lineage-specific tissues [14–16]. However, it is unclear if the transcriptional heterogeneity of planarian stem cells is itself required for some or all of their unique properties, as the molecular mechanisms that mediate this transcriptional heterogeneity remain largely unknown.

One category of proteins with strong potential to act as regulators of genome stability and transcriptional dynamics in planarian stem cells is chromatin modifying proteins. This group includes enzymes that add covalent modifications to histone proteins, those that remove them, proteins that bind specifically modified histones, and complexes that open, close, and/or remodel the chromatin structure [17–19]. These different modifiers work together as part of multilayered, synergistic mechanisms that provide increased specificity and robustness to gene regulatory networks. For example, histone H3 lysine 4 trimethylation (H3K4me3), a modification that correlates strongly with gene expression in many different organisms [11, 20–22], is added by specific lysine methyltransferases (Set1A/B and MLL1-4, or KMT2A-D), removed by specific lysine demethylases (KDM5A-D), and recognized by specialized motifs in chromatin “reader” proteins e.g., BPTF [23, 24]. BPTF is the largest subunit of the Nucleosome Remodeling Factor (NuRF), a complex that executes its function by shifting nucleosomes bidirectionally in 10bp increments to make the underlying genomic sequence more accessible [25–27]. In addition, knockdown of bptf in Xenopus embryos phenocopies the axial, blood, and gut defects seen in embryos with reduced H3K4me3 (via wdr5 knockdown) [24]. Together, these data support the hypothesis that H3K4me3 is translated into transcriptional and biological function by NuRF.

Despite the substantial data supporting the H3K4me3 writer-reader mechanism, many questions remain about the functional link between H3K4me3 and transcriptional activation [28, 29]. Puzzlingly, although H3K4me3 shows a strong, positive correlation with gene expression across most gene loci and in many different organisms [11, 20–22], loss of H3K4me3 does not consistently lead to loss of transcription [22, 28, 30, 31]. This suggests that for many gene loci H3K4me3 may be a symptom, not a cause, of active transcription. Yet the null mutation of any of the H3K4me3 KMTases identified in mice and *Drosophila* causes embryonic lethality [32–36]. In other words, despite sharing a common enzymatic activity and substrate, H3K4me3 KMTases each have essential in vivo functions and do not compensate for each other during early development. Does this imply that each enzyme targets different gene loci? Do they each create distinctive H3K4me3 patterns that are translated differently by downstream binding proteins? Are the essential functions of these KMTases even mediated through the methylation of histones?

Planarians are an excellent model for addressing these questions. First, as in other animals, conserved H3K4me3 KMTases have non-redundant in vivo functions in *Schmidtea mediterranea* (*S.med*). Knockdown of *Smed-set1* causes progressive stem cell dysfunction, tissue regression, and death, whereas knockdown of *Smed-mll1/2* induces motility defects due to the gradual loss of epidermal cilia [22, 37]. Second, these distinct phenotypes are clearly linked to the specific genomic targets of Smed-Set1 versus Smed-MLL1/2 in planarian stem cells: Set1 methylates genes with known stem cell functions, while MLL1/2 methylates highly conserved cilia genes [22]. In addition, these enzymes create significantly different methylation patterns at their respective genomic targets: Set1-dependent H3K4me3 peaks are significantly wider than an average H3K4me3 peak, but MLL1/2-dependent peaks are significantly narrower [22]. Together, these data not only helped to answer some of the above questions (as individual planarian H3K4me3 KMTases do target distinct and phenotypically-relevant genes), but also provided a clear and tractable model for addressing the next logical question: does BPTF translate the distinct H3K4me3 patterns created by Set1 and MLL1/2 into functional genomic output?

In this study, we test the hypothesis that the local concentration of BPTF, and thus NuRF activity, is higher at Set1 target genes due to their wider H3K4me3 peaks. We used multiple genomic assays to identify changes in the chromatin state and transcriptional output of planarian stem cells after *bptf* knockdown. We find that BPTF is essential for both chromatin and transcriptional stability in planarian stem cells, as loss of its activity not only causes significant changes in chromatin accessibility at specific loci but also targets genes that themselves are essential for maintenance of a stable chromatin landscape and gene regulatory networks. Many of these BPTF target genes are shared with those of Set1, but not those of MLL1/2, a finding that was further validated by in vivo functional data showing that knockdown of *bptf* mimics the stem cell deficiency phenotype seen after knockdown of *set1.* Collectively, these results provide new insight into both the genome regulation of planarian stem cells and the mechanism linking H3K4me3 to functional changes in gene expression.

## RESULTS

### The genome of *Schmidtea mediterranea* encodes a homolog of NuRF protein BPTF

To test the hypothesis that NuRF mediates functional differences in gene expression at Set1 versus MLL1/2 target genes in planarian stem cells (**Figure 1A**), we first identified the likely homolog of NuRF subunit BPTF in *S.med* using a reciprocal BLAST [38] strategy. This search identified a single gene encoding a protein with 27% sequence identity to human BPTF and 28% identity to the Drosophila homolog NURF301 (calculated using Clustal Omega; in the same matrix DmNURF301 has 34.5% identity with human BPTF). The predicted protein domains in this *S.med* homolog are highly conserved with those of the human and Drosophila BPTF proteins (**Figure 1B**). Moreover, when we examined an alignment of the PHD2 motif of these species more closely (**Figure 1C**), we saw that all four aromatic residues essential for H3K4me3 binding [39] are conserved in *S.med* BPTF (asterisks).

**Figure 1:**
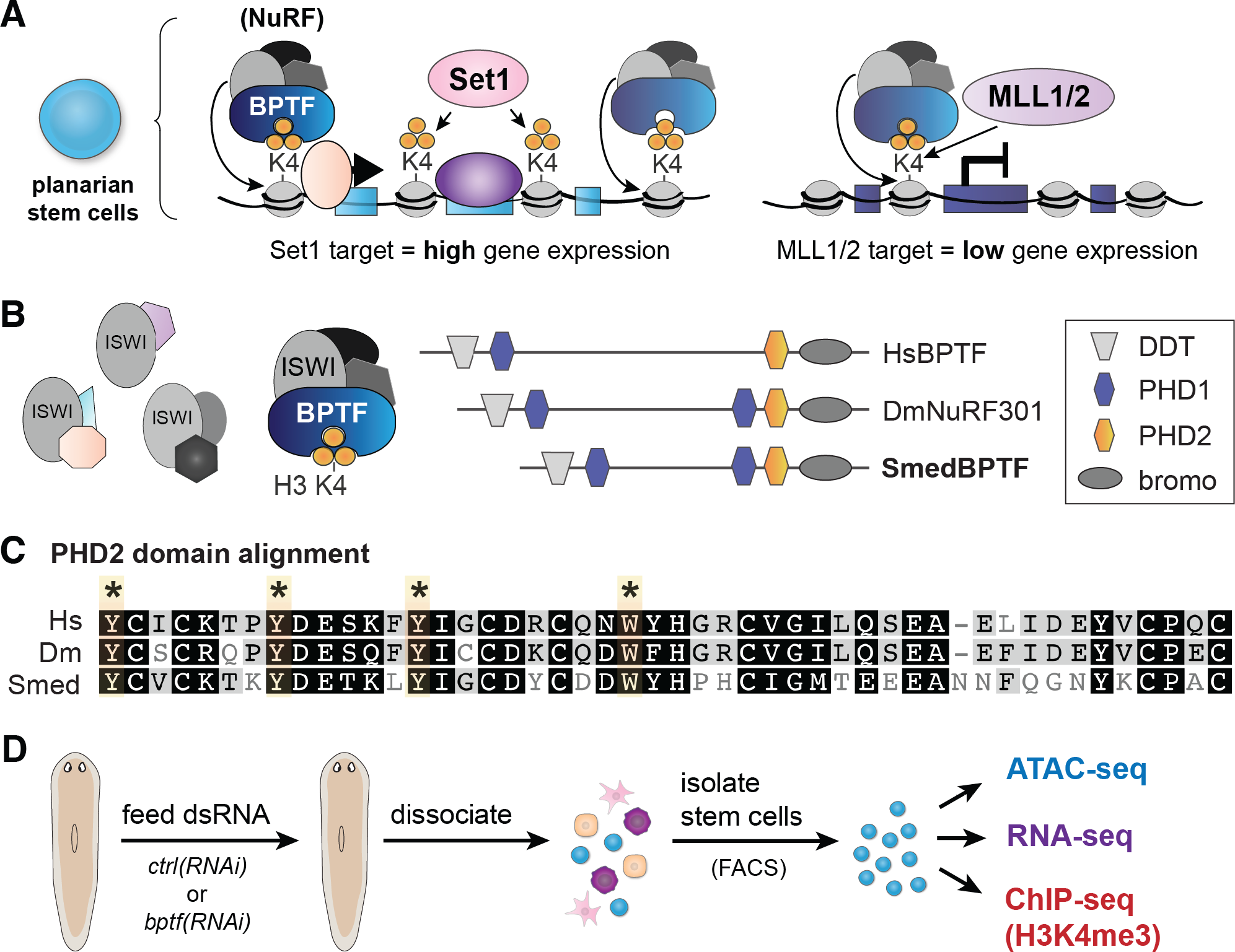
The genome of *Schmidtea mediterranea* encodes a homolog of BPTF, the H3K4me3- binding subunit of the Nucleosome Remodeling Factor (NuRF) complex. A) Model of proposed mechanism through which broad peaks of histone H3 lysine 4 trimethylation (H3K4me3) catalyzed by Set1 result in active transcription of its target gene loci but the narrow H3K4me3 peaks catalyzed by MLL1/2 are less active. NuRF = Nucleosome Remodeling Factor complex. Orange circles represent methyl groups (in set of three = trimethylation), peach and purple ovals represent parts of the transcription machinery. Blue boxes represent exons of target genes. B) Illustration comparing the domain structures of human BPTF, Drosophila NURF301 and a BPTF/NURF homolog identified in the planarian species *Schmidtea mediterranea* (*S.med*). C) Alignment of the PHD2 domain from human (NP_872579.2), *Drosophila*, and *S.med* BPTF. Those amino acids known to be critical for its H3K4me3-binding [91] are highlighted and asterisked. D) Schematic showing the strategy used to assess the functional, in vivo impact of BPTF on the chromatin state and genome output of planarian stem cells.

Having identified a highly conserved homolog of BPTF, we then examined its functional effects on genome organization and transcriptional output in planarian stem cells. After cloning *S.med* bptf and generating dsRNA from its sequence, we fed this dsRNA to adult planarians to trigger RNA interference (RNAi) and deplete matching endogenous *bptf* transcripts (**Figure 1D**). In parallel, we fed separate animals with dsRNA for the *C. elegans* unc-22 gene, as its sequence is not found in the planarian genome and it serves as a control for changes induced by the RNAi process itself [40]. After administering three or four doses of dsRNA, we dissociated the treated animals into single cell suspensions, stained them with Hoechst nuclear dye, and isolated the X1 population of stem cells (S/G2/M cells with >2n DNA) by flow cytometry as previously reported [41]. We collected multiple replicates of stem cells per *control(RNAi)* and *bptf(RNAi)* and subjected them, separately, to ATAC-seq, RNA-seq, and ChIP-seq (**Figure 1D**).

### Knockdown of *bptf* leads to loss of chromatin accessibility at gene promoters

As BPTF is the largest subunit of the NuRF chromatin remodeling complex [25], we predicted that knockdown of its planarian homolog would cause loss of chromatin accessibility at NuRF target genes. We tested this prediction by performing ATAC-seq [42] on isolated X1 stem cells (**Figure 2A, Supplemental Figure S1A**) and aligning the data to the newest *S.med* genome assembly (S3h1) [43]. We detected thousands of ATAC-seq peaks in both samples, including those with measurable differences between *control(RNAi)* and *bptf(RNAi)* and those with no significant change (**Figure 2B**). After mapping these peaks to the nearest annotated gene model, we found that most ATAC-seq peaks in *control(RNAi)* stem cells map near promoters and a significant number map to distal intergenic regions (**Figure 2C, Supplemental Figure S1B**). We then used csaw [44] to identify those peaks with significant changes in ATAC-seq signal in *bptf(RNAi)* stem cells. More than half of the differentially-accessible (DA) peaks we detected mapped to gene promoters, with a relatively smaller fraction mapping to distal intergenic regions (**Figure 2D; Supplemental Figure S1C**). The majority of these DA peaks lose ATAC-seq signal in *bptf(RNAi)* stem cells (**Figure 2E, Supplemental Figure S1C**), which is expected given that BPTF is part of the NuRF remodeling complex. These differences are also unambiguously observed when we averaged ATAC-seq signal across all annotated gene models: ATAC-seq signal from *bptf(RNAi)* stem cells was significantly lower at gene promoters (i.e., +/- 1kb of the TSS; **Figure 2F**), but not detectably changed at distal intergenic loci (**Figure 2G**). These data support the prediction that SmedBPTF is a conserved subunit of the NuRF complex and reveal that BPTF largely recruits NuRF activity to gene promoters in planarian X1 stem cells.

**Figure 2:**
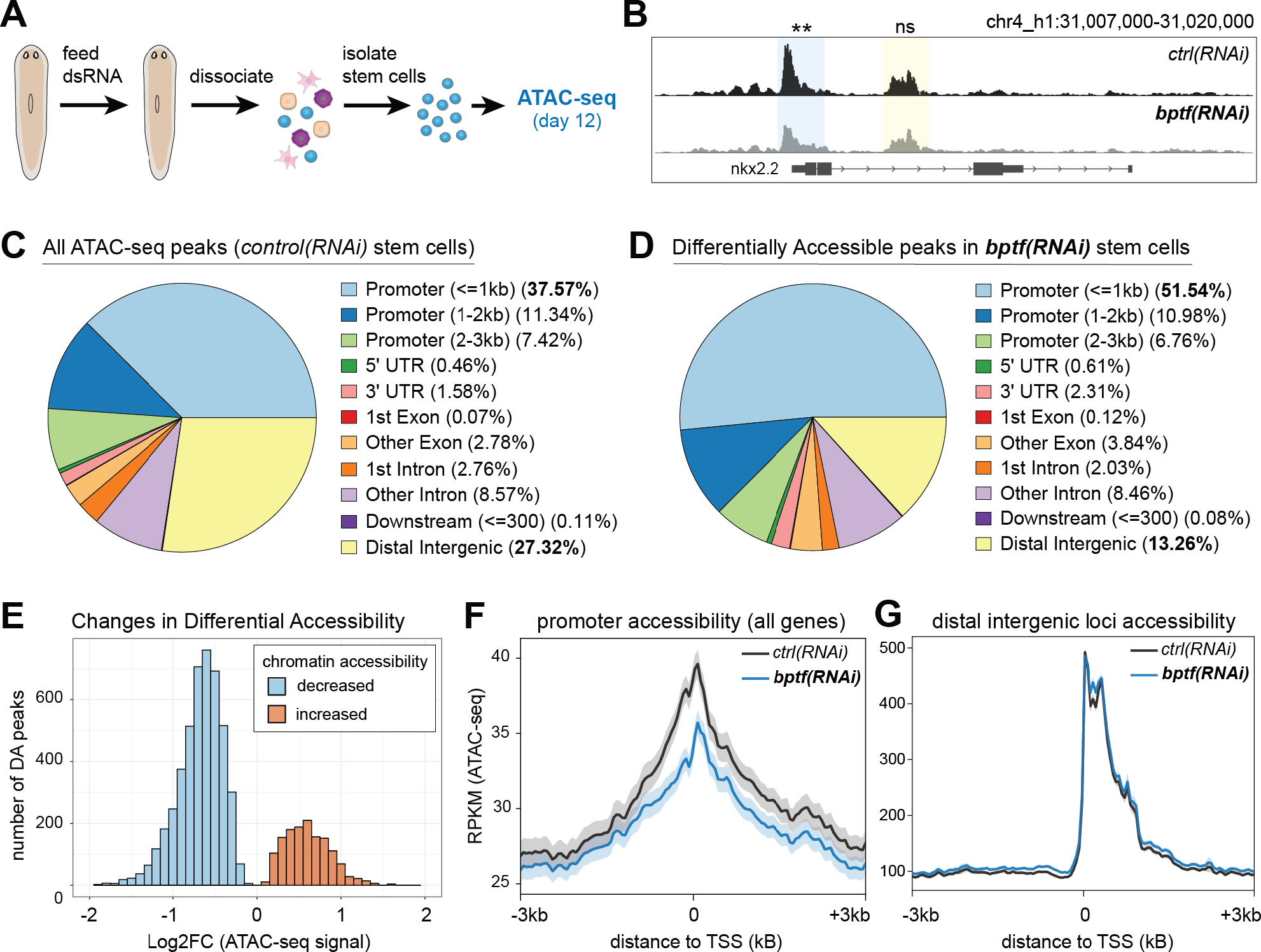
Knockdown of *bptf* leads to loss of chromatin accessibility at gene promoters in planarian stem cells. A) Schematic of the experimental setup used to assay the chromatin state of planarian stem cells isolated from *control(RNAi)* and *bptf(RNAi)* animals. B) A representative locus at which there are multiple ATAC-seq peaks, including one that shows significant loss of accessibility after *bptf(RNAi)*. Gene nkx2.2 = h1SMcG0021724. C) Pie chart summarizing the locations of ATAC-seq peaks across the planarian genome (relative to their nearest gene models) in *control(RNAi)* stem cells. D) Pie chart summarizing the locations of differentially accessible (DA) peaks in *bptf(RNAi)* stem cells (compared to *control(RNAi)* stem cells). E) Histogram summarizing the log2FC of all differentially accessible (DA) ATAC-seq peaks in *bptf(RNAi)* stem cells (binwidth = 0.1 log2FC). 4652 peaks have decreased accessibility (blue bars), 1411 have increased accessibility (orange bars). F) Profile plot comparing ATAC-seq signal in *control(RNAi)* and *bptf(RNAi)* stem cells averaged across all genes. Signal is represented as Reads Per Kilobase per Million mapped reads (RPKM). The shaded area shows the 95% confidence interval for each condition. G) Profile plot comparing ATAC-seq signal at distal intergenic peaks (identified in C).

### Knockdown of *bptf* leads to significant changes in gene expression

As changes in chromatin accessibility have the potential to affect transcription, we then asked if knockdown of *bptf* also impacted gene expression in planarian stem cells. Again, we isolated X1 stem cells by flow cytometry from *control(RNAi)* and *bptf(RNAi)* animals, in this case to be used for total RNA isolation and RNA-sequencing (**Figure 3A**). We isolated stem cells at two time points post-RNAi treatment to assess the timing and consistency in gene expression changes. After isolating total RNA from these cells and performing RNA-sequencing, we detected many significant changes at both time points (**Figure 3B**). Overall, the changes in gene expression, both up and down-regulated, were consistent between time points (**Figure 3C, Supplemental Figure S2A**). In addition, when we grouped genes into four broad expression categories based on average TPM (transcripts per million) in *control(RNAi)* stem cells, we did find an overall correlation between average ATAC-seq signal at promoters and average gene expression (**Supplemental Figure S2B**). Interestingly, we saw a relatively even distribution of up and down-regulated genes in *bptf(RNAi)* stem cells (**Figure 3B**), which was unexpected given the overall loss of chromatin accessibility at gene promoters in these cells (**Figure 2E-F)**. Moreover, when we compared changes in chromatin accessibility (ATAC-seq for each identified DA peak) with changes in gene expression (RNA-seq for the nearest mapped gene), there was no correlation (**Figure 3D**). This was also true when we restricted the analysis to DA peaks at promoters (**Supplemental Figure S2C**).

**Figure 3:**
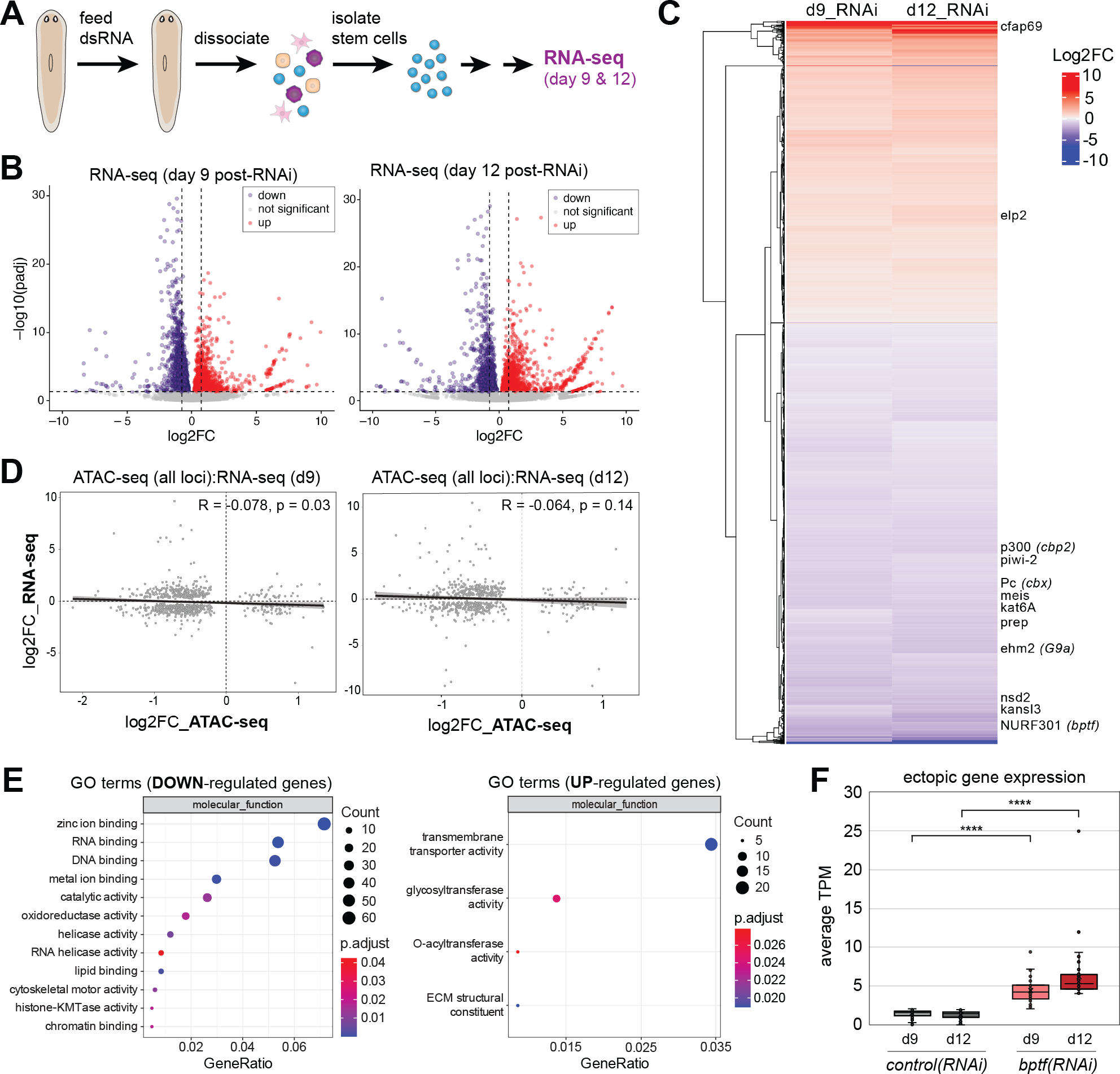
Loss of BPTF leads to significant changes in gene expression that indicate the dysregulation of transcription and chromatin regulation. A) Schematic of the experimental setup used to assay changes in gene expression in planarian stem cells due to *bptf* knockdown. B) Volcano plots showing transcript changes in *bptf(RNAi)* stem cells versus *control(RNAi)* as detected by RNA-seq. Left plot shows data from stem cells isolated 9 days post-RNAi; right plot shows data from 12 days post-RNAi. Blue dots = significantly down-regulated (pAdj < 0.05), orange dots = significantly up-regulated (pAdj < 0.05). Vertical dashed lines = log2FC +/-0.75. Horizontal dashed line = linear pAdj 0.05. C) Clustered heat map of day 9 and day 12 RNA-seq data in B; only genes with significant differential expression (DE; pAdj <0.05) at both time points are included. DE between time points correlates strongly (see **Supplemental Figure S2A**) D) Correlation plot comparing changes in accessibility of all DA peaks (Figure 2) with changes in expression (RNA-seq) of their nearest gene. Left plot shows correlation with day 9 RNA-seq data and right plot shows data from day 12 RNA-seq. Lack of statistical correlation shown by R (Pearson correlation coefficient) and p-values. E) GO Term enrichment analysis of genes down-regulated after *bptf* knockdown (left plot) or up- regulated (right plot). F) Box plot showing the average TPM (transcripts per million) for genes significantly up-regulated in *bptf(RNAi)* stem cells but that have little to no expression (<2 TPM) in control stem cells (**** = p- value < 0.0001). Many of these genes are normally restricted to differentiated cells, such as neural, muscle, or intestinal cells, according to published single cell RNA-seq ATLASes [12, 13].

The discrepancy between the broad correlation of chromatin accessibility with gene expression in control stem cells and the lack of correlation at *bptf(RNAi)* sensitive loci was at first puzzling. We surmised that loss of BPTF (and its recruited NuRF activity) may significantly impact the expression of transcriptional regulators themselves, causing additional, indirect changes in gene expression. To assess the validity of this hypothesis, we performed a Gene Ontology (GO) enrichment analysis on the genes we identified as down-regulated in *bptf(RNAi)* stem cells and found that terms associated with transcriptional regulation were significantly enriched (**Figure 3E**). Moreover, genes encoding proteins associated with both chromatin activation (p300/cbp2, kat6A) and chromatin silencing (Polycomb/cbx, ehm2/g9a) were down-regulated in *bptf(RNAi)* stem cells (**Figure 3C**), suggesting that loss of BPTF may lead to secondary increases in gene expression by reducing the expression of chromatin silencing machinery. To assess this possibility further, we compared the absolute expression (in transcripts per million, or TPM) of genes we detected as up-regulated and found many genes that were ectopically expressed in *bptf(RNAi)* stem cells i.e., negligible expression in *control(RNAi)* stem cells but significant expression in *bptf(RNAi)* stem cells (**Figure 3F**). Together, these data suggest that BPTF regulates transcription both directly and indirectly.

### BPTF-mediated accessibility is enhanced at promoters with H3K4me3

Because knockdown of *bptf* caused significant changes in the expression of many other chromatin modifiers, we first sought to determine whether H3K4me3 itself was affected by loss of *bptf.* We isolated X1 stem cells from both *control(RNAi)* and *bptf(RNAi)* animals and performed ChIP-seq with an antibody to H3K4me3 (**Figure 4A**). As shown previously in *S.med* [22, 43] and many other organisms [20, 45, 46], nearly all MACS2-called H3K4me3 ChIP-seq peaks were detected at gene promoters (**Figure 4B-C**). Moreover, we did not observe a detectable change in H3K4me3 after knockdown of *bptf* (**Figure 4B**). We then asked how many gene promoters have both an H3K4me3 peak and an ATAC-seq peak in *control(RNAi)* stem cells (**Figure 4D**). We found that 28% of all promoters (+/- 1 kb) with an ATAC-seq peak also had an H3K4me3 peak within 1kb. Similarly, a quarter of all promoters with an H3K4me3 peak also had an ATAC-seq peak.

**Figure 4:**
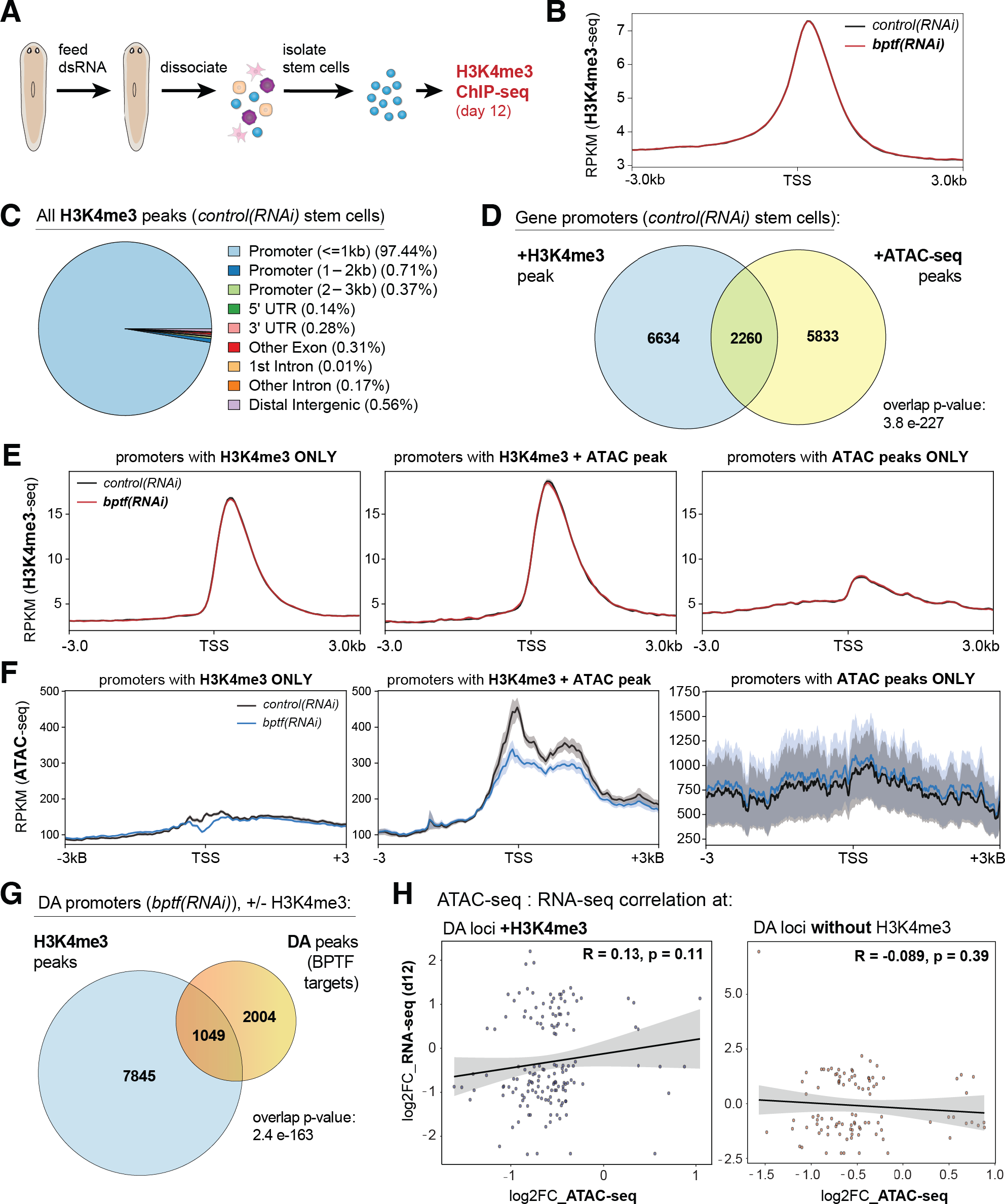
BPTF has a stronger effect on transcription at promoters with H3K4me3. A) Schematic of the experimental setup used to assay H3K4me3 across the planarian genome of stem cells isolated from *control(RNAi)* and *bptf(RNAi)* animals. B) Profile plot comparing H3K4me3 signal in *control(RNAi)* and *bptf(RNAi)* stem cells averaged across all genes. C) ChIPseeker plot showing the distribution of mapped H3K4me3 peaks (MACS2) in *control(RNAi)* stem cells. D) Venn diagram showing the number of promoters in planarian stem cells with an H3K4me3 peak, an ATAC-seq peak, or both. The p-value for the overlap was calculated using the hypergeometric distribution. E) Profile plot comparing H3K4me3 signal in *control(RNAi)* and *bptf(RNAi)* stem cells at the gene promoters in each group of the Venn in D (H3K4me3 only, H3K4me3 + ATAC peak, and ATAC peak only). F) Profile plot comparing ATAC-seq signal in *control(RNAi)* and *bptf(RNAi)* stem cells for gene promoters in each group of the Venn shown in D. G) Venn diagram showing the overlap between gene promoters with an H3K4me3 peak (MACS2) and those with a Differentially Accessible (DA) peak in *bptf(RNAi)* stem cells. H) Plots correlating changes in chromatin accessibility (log2FC ATAC-seq) with changes in transcription (log2FC RNA-seq) for those genes with an H3K4me3 peak at promoter (left plot) and those without one (right plot). Although the R value (Pearson correlation coefficient) for the correlation of ATAC-seq and RNA-seq changes at genes with H3K4me3 promoter peaks is not significant, it is positive and larger than that of the correlations in Figure 3D. Those genes without H3K4me3 at their promoters show a negative correlation.

To determine whether the presence of an H3K4me3 peak impacts BPTF-mediated chromatin accessibility, we examined the average H3K4me3 (**Figure 4E**) and ATAC-seq signal (**Figure 4F**) at those genes annotated with: 1) an H3K4me3 peak only; 2) both an H3K4me3 peak and an ATAC-seq peak; and 3) an ATAC-seq peak only (groups based on **Figure 4D**). The analysis revealed that *bptf(RNAi)* has the largest effect at those genes with both H3K4me3 and ATAC-seq peaks (**Figure 4F, middle**). Genes with “ATAC-seq only” promoters have relatively higher levels of ATAC-seq signal overall (**Figure 4F, right**), but this signal is highly variable between genes in this group (as shown by the wide 95% confidence intervals). We noted that this third group includes significantly more “unknown” gene models than the promoters marked by H3K4me3 and wondered whether the chromatin state at these “unknown” genes was different from that of “high confidence” models (defined in [43]). However, when we divided the genes in the “ATAC-seq only” group into “high confidence” models versus “rest”, we saw no detectable difference in overall signal or on the effect of *bptf(RNAi)* (**Supplemental Figure S3A**).

After observing that the chromatin accessibility of genes with both H3K4me3 and ATAC-seq peaks at their promoters were more sensitive to loss of BPTF, we then asked if these genes also experienced a greater impact on their transcription. We split DA (differentially-accessible) ATAC-seq promoter peaks into two groups based on their chromatin state (**Figure 4G**): 1049 peaks had H3K4me3 peaks at their promoters (DA loci +H3K4me3) and 2004 did not have an H3K4me3 peak (DA loci -H3K4me3). We then asked if the changes in accessibility of the DA peaks in each group correlate with respective changes in gene expression (**Figure 4H**). Although neither group showed a statistically significant correlation (R and p values), those genes with both an H3K4me3 peak at their promoter and a BPTF-dependent change in ATAC-seq did show a positive, though not significant, correlation with RNA-seq changes. On the other hand, those genes without H3K4me3 at their promoters trended toward a negative correlation. This was true when comparing to RNA-seq changes at day 12 post-RNAi (**Figure 4H**) and day 9 post-RNAi (**Supplemental Figure S3B**). These data show that H3K4me3 distinguishes BPTF-dependent loci in the genomes, yet also suggests that not all H3K4me3 loci are regulated by the same mechanisms.

### Set1 targets are significantly and functionally regulated by NuRF

After determining that genes marked by H3K4me3 are more sensitive to BPTF-dependent chromatin remodeling, we returned to our original hypothesis: that the broad peaks of H3K4me3 catalyzed by Set1 recruit a higher local concentration of NuRF complex than MLL1/2 targeted genes, leading to greater genome accessibility and increased transcriptional activity (**Figure 1A**). After realigning our published H3K4me3-ChIP-seq data [22] to the S3h1 genome assembly [43], we confirmed that loci targeted by Set1 are significantly wider (on average) than those targeted by MLL1/2 in planarian stem cells (**Figure 5A; Supplemental Figure S4A**). We then compared the ATAC-seq signal at Set1 and MLL1/2 target gene loci. As predicted by our model, Set1 target genes have significantly more ATAC-seq signal than MLL1/2 target genes (**Figure 5B-C**). Notably, both Set1 and MLL1/2 target genes lose a significant amount of ATAC-seq in *bptf(RNAi)* stem cells (**Figure 5C**). This finding supports our model, as BPTF is not predicted to discriminate between gene loci but simply bind H3K4me3 where it exists.

**Figure 5.**
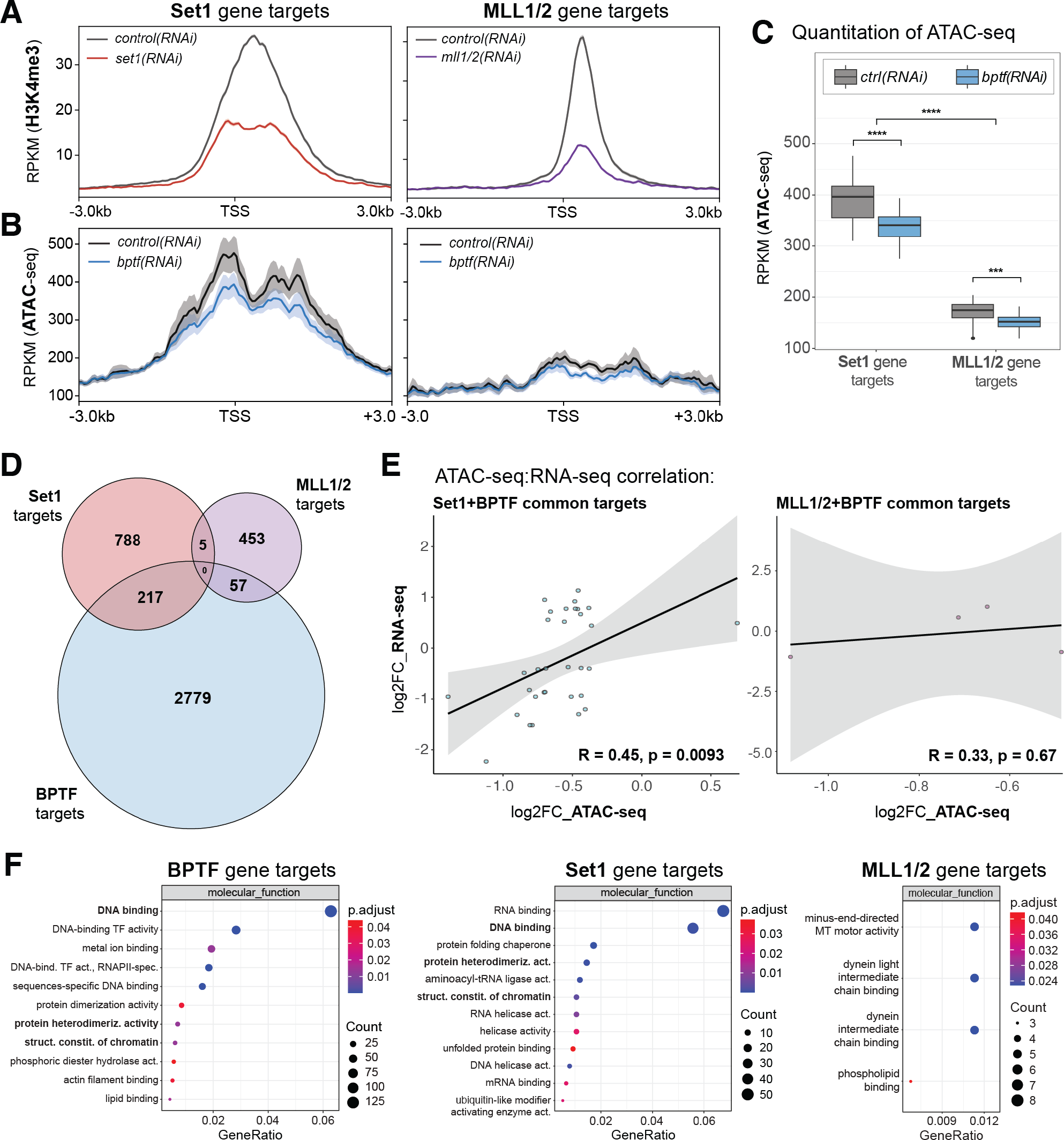
BPTF shares functional genomic targets with Set1. A) Profile plots averaging H3K4me3 ChIP-seq (from [22]) across all genes mapped to the newest genome assembly [43]. Left plot = H3K4me3 ChIP-seq data from *set1(RNAi)* and matched *control(RNAi)* stem cells, right plot = H3K4me3 ChIP-seq data from *mll1/2(RNAi)* and matched *control(RNAi)* stem cells. B) Profile plot comparing ATAC-seq data from *control(RNAi)* and *bptf(RNAi)* stem cells at Set1 (left) and MLL1/2 (right) gene targets. C) Quantitation and statistical analysis of ATAC-seq data in B; statistical significance was determined using a Wilcoxon test (p-value with Bonferroni correction). * = p≤ 0.05, ** = p≤ 0.01, *** = p≤ 0.001, **** = p≤ 0.0001). D) Venn diagram comparing genes identified as likely targets of Set1, MLL1/2, and BPTF (i.e., genes with significant loss of ATAC-seq in *bptf(RNAi)* stem cells). p-val = 8.65e-74 for BPTF targets overlap with Set1 targets; p-val= 1.04e-75 for BPTF targets overlap with MLL1/2. E) Plots correlating significant changes in chromatin accessibility (ATAC-seq) with significant changes in gene expression (RNA-seq, day 12) in *bptf(RNAi)* stem cells. Left plot = genes targeted by Set1 and BPTF, right plot = genes targeted by MLL1/2 and BPTF (from D). The R and p-values are displayed in each plot. F) GO Term enrichment analysis for each group of genes in the Venn diagram in D.

In line with these results, when we compare Set1 target genes with those of MLL1/2 and BPTF (i.e., promoters with differentially-accessibly ATAC-seq peaks), we find that more BPTF target genes overlap with Set1 targets versus MLL1/2 targets (21% of Set1 targets overlap with BPTF-dependent DA peaks compared to 11% of MLL1/2 targets; **Figure 5D**). Moreover, when we correlated ATAC-seq changes with RNA-seq changes at Set1 target genes, we saw a significant, positive correlation (**Figure 5E, left plot**). In contrast, MLL1/2 target genes with BPTF sensitive ATAC-seq peaks did not show a significant correlation with changes in RNA-seq (**Figure 5E, right plot**). In addition, Gene Ontology (GO) enrichment analysis reveals that Set1 targets and BPTF targets share many common molecular functions, including DNA binding (GO:0003677), whereas the MLL1/2 targets are distinct in their classifications (**Figure 5F**). These data strongly support our hypothesis that BPTF mediates the different outputs of Set1 and MLL1/2 target genes (**Figure 1A**).

We then wondered if BPTF/NuRF activity was linked to the different H3K4me3 peak widths its targets (**Supplemental Figure S4A**), irrespective of the specific KMTase adding them. We addressed this question by dividing all H3K4me3 peaks in *control(RNAi)* stem cells into three categories based on their width: “narrow”, “medium”, and “broad” (**Supplemental Figure S4B-D**). After plotting the average ATAC-seq signal in each group (**Supplemental Figure S4E**), we found the overall ATAC-seq signal increased with H3K4me3 peak width (**Supplemental Figure 4E-G**). However, several details in these data suggest that Set1 and MLL1/2 target genes are distinct subsets of their peak width groups.

First, when comparing all “narrow”, “medium”, and “broad” H3K4me3 peaks, regardless of the responsible KMTase, there is a correlating increase in H3K4me3 peak amplitude (**Supplemental Figure S4D**). However, Set1 and MLL1/2 target genes have comparable H3K4me3 peak amplitudes, on average, despite significantly different widths (**Figure 5A**). In addition, when we quantitate BPTF-dependent changes in ATAC-seq at “broad” H3K4me3 peaks, it does not reach the threshold of statistical significance (**Supplemental Figure S4F-G**). In contrast, Set1 target genes have similarly broad H3K4me3 peaks (**Supplemental Figure S4A, C**) but do show statistically significant changes in ATAC-seq in *bptf(RNAi)* stem cells (**Figure 5C**). Second, MLL1/2 targets have relatively low chromatin accessibility given their H3K4me3 peak width and amplitude (**Figure 5A-B, Supplemental Figure S4A, C-E**). These findings suggest that although H3K4me3 peak width is a relevant feature of BPTF-dependent loci, there are likely additional contexts at Set1 loci that enhance BPTF function further.

Although the presence of Set1-mediated H3K4me3 at gene promoters did significantly enhance the overall effects of BPTF/NuRF activity, we also observed many individual loci at which there was a BPTF-dependent loss of chromatin accessibility and a correlating loss of gene expression without detectable H3K4me3 at the gene promoter (**Figure 4G-HC**). Notably, many of these loci have multiple BPTF-dependent ATAC-seq peaks mapped to their genes (**Supplemental Figure S5; Supplemental Table S1**), including those encoding the chromatin silencing proteins Polycomb (or cbx in mammals) and emh2 (or G9a) (**Supplemental Figure S5**). Collectively, these “multi-DA-peak” loci did not show a significant correlation with BPTF dependent changes in gene expression. Yet they suggest a potential mechanism underlying distinctions between *bptf(RNAi)* and *set1(RNAi)* stem cells.

### Knockdown of chromatin remodeling protein BPTF mimics knockdown of KMTase Set1

Together, these genomic data strongly support the hypothesis that BPTF is particularly important at Set1 target genes. To test if this extends to its in vivo function, we fed adult planarians with dsRNA matching bptf, set1, mll1/2, and control genes. As shown in previous studies [22, 37], *set1(RNAi)* animals showed signs of tissue regression, including progressive head regression (**Figure 6A**), within two weeks of dsRNA treatment and were all dead within three weeks. In contrast, *mll1/2(RNAi)* worms did not show signs of head regression even after several weeks but did develop motility defects due to loss of ventral cilia [37, 47]. RNAi of *bptf* caused a penetrant and morphologically similar phenotype to that of *set1*, although its progression developed slower than the *set1(RNAi)* phenotype (**Figure 6A**). Importantly, because the lethal phenotypes of *set1(RNAi)* and *bptf(RNAi)* are relatively quick (2-3 weeks) compared to the slower onset of the *mll1/2(RNAi)* motility defect (3-4 weeks), we cannot conclude from these phenotypes that *S.med* BPTF only mediates changes in Set1 target gene expression.

**Figure 6.**
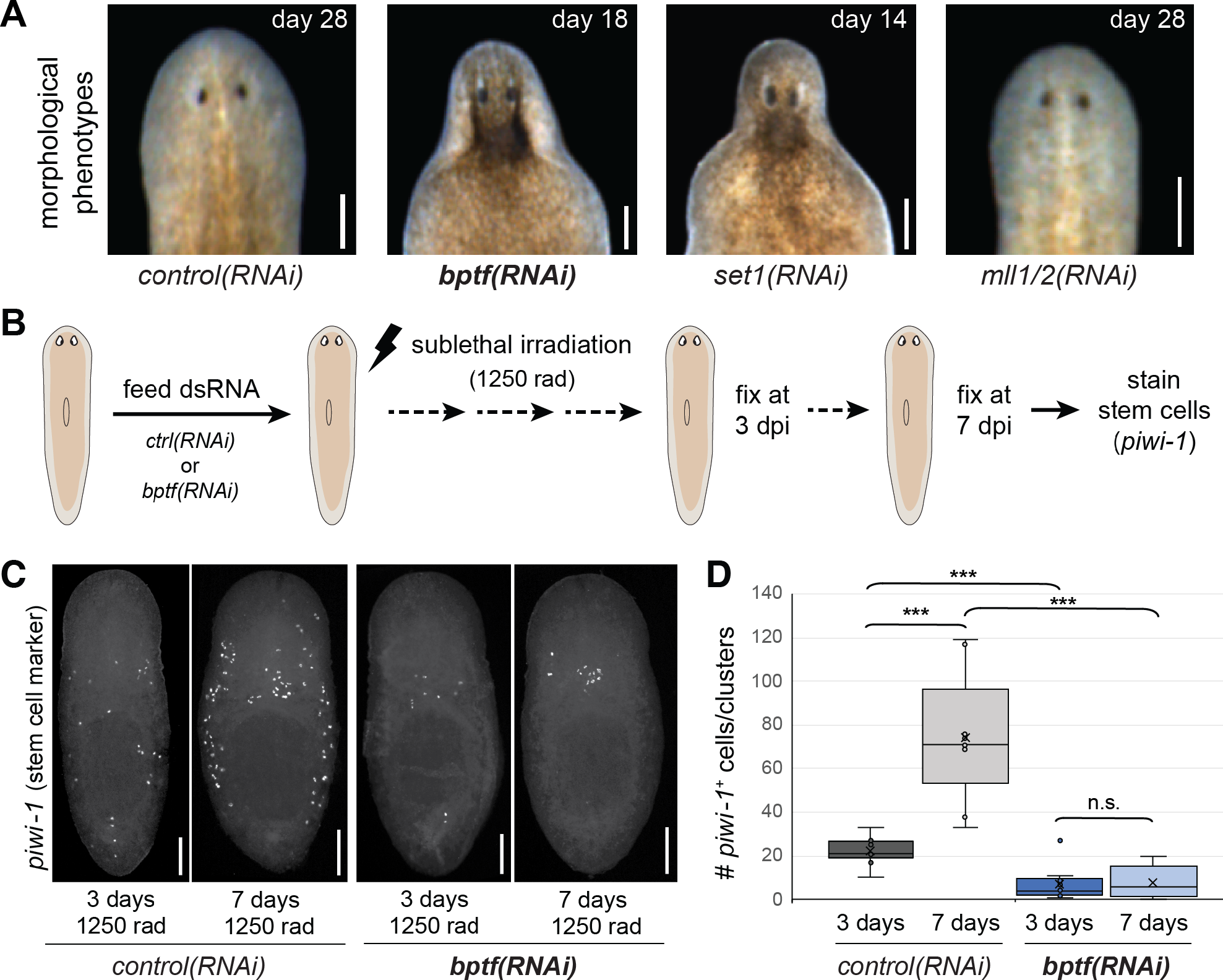
RNAi knockdown of *Smed-bptf* phenocopies that of *Smed-set1*, but not *Smed-mll1/2*. A) Live images showing the morphological phenotypes of *bptf(RNAi)* planarians (*S.med*) compared to *control(RNAi)*, *set1(RNAi)*, and *mll1/2(RNAi)* animals. Scale bars = 200µm. B) Schematic of the experimental setup used to test planarian stem cell function in RNAi worms. In normal conditions, a small but significant number of planarian stem cells will survive 1250 rad (12.5 Gy) γ-radiation (3 days post-irradiation, or dpi), then resume proliferating (7 dpi) to restore the population [92]. C) Representative Whole mount In situ Hybridization (WISH) images of *control(RNAi)* and *bptf(RNAi)* planarians stained with the stem cell marker *piwi-1*. Scale bars = 250µm. D) Quantitation of *piwi-1^+^* cells per animal for all animals included in E. *** = p-value < 0.001.

In planarians, head and tissue regression can be caused by the failure of various different cellular processes, including the maintenance and differentiation of their stem cell population.

Previous studies from our lab and others [22, 37, 48] have shown that *set1(RNAi)* worms show defects in stem cell function, including an inability to recover from population-depleting doses of radiation. To determine how knockdown of *bptf* affects stem cell function, we treated *control(RNAi)* and *bptf(RNAi)* worms with 1250 rads (12.5Gy) ionizing radiation and then stained subsets of these worms at 3 and 7 days post radiation with a marker for planarian stem cells (**Figure 6B**). As expected, *control(RNAi)* worms showed an initial depletion of most, but not all, *piwi-1+* stem cells followed by significant restoration of this population by day 7 (**Figure 6C, D**). In contrast, *bptf(RNAi)* worms had significantly fewer stem cells at 3 days post-radiation treatment and failed to show significant recovery at day 7 (**Figure 6D**). Notably, this functional effect is also seen in the stem cells of *set1(RNAi)* worms [22], supporting the hypothesis that the phenotypic effects of *set1(RNAi)* and *bptf(RNAi)* share common cellular origins.

## DISCUSSION

Planarian stem cells have many fascinating cellular characteristics and are required for two extraordinary features of their organism: 1) planarians can repeatedly and reproducibly regenerate complex tissues and organs, and 2) they are effectively immortal. Although many genes that are important for the function of the stem cells have been identified, the mechanisms regulating their expression are poorly understood. Here, we examined the role of the chromatin remodeling complex NuRF in planarian stem cell biology. By knocking down its largest and specific subunit, BPTF, we both targeted the NuRF component that provides its chromatin-state specificity and tested the hypothesis that local concentrations of NuRF are relevant to its effects on transcription in vivo [24, 49]. Our findings show that BPTF, and the remodeling activity it recruits to its gene loci, are critical for planarian stem cell function (**Figure 7**).

**Figure 7.**
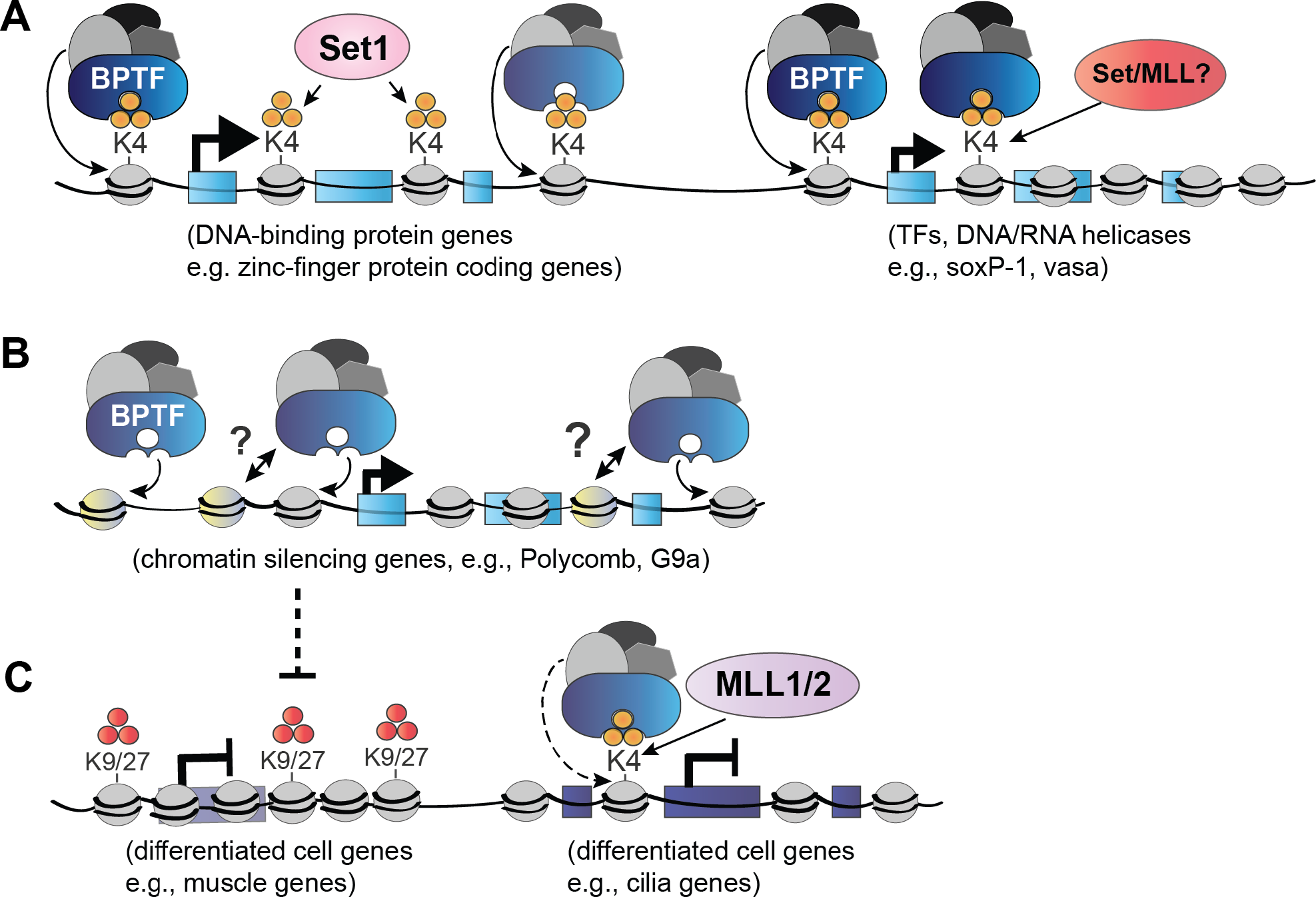
BPTF regulates transcription in multiple ways. Model summarizing findings and potential implications of the data reported in this paper. A) BPTF-mediated NuRF activity is concentrated at the promoters of genes that are also marked by H3K4me3, particularly Set1 target genes with wide H3K4me3 peaks. B) A smaller subset of BPTF/NuRF affected promoters do not show enrichment of H3K4me3 but do correlate with changes in gene expression. Often these genes have multiple affected peaks of chromatin accessibility mapped to their promoters, suggesting there may be other chromatin features that recruit BPTF to these loci. C) Some genes marked by H3K4me3, including MLL1/2 target genes, remain closed and transcriptionally inactive. Data reported here suggests some of these genes may be indirectly regulated by BPTF/NuRF, as knockdown of *bptf* reduces the expression of genes that encode chromatin silencing proteins.

We set out to test a simple hypothesis: that the broad peaks of H3K4me3 at Set1 target genes recruits a higher molar ratio of NuRF (via BPTF), which then opens the chromatin at these genes and mediates their relatively high expression. This hypothesis is not only relevant to question of planarian biology, as differences in H3K4me3 peak width are also observed in many other organisms, including mammals [50, 51]. Further, wide H3K4me3 peaks have been shown to mark particular loci, such as tumor suppressor genes [50]. Our data largely supports our mechanistic model (**Figure 1A**). Using ATAC-seq, we identified many specific regions of the planarian genome at which chromatin accessibility changed significantly after *bptf* knockdown. We confirmed that, as expected, the chromatin at Set1 target genes is significantly more accessible than that at MLL1/2 targets. Moreover, genes with both a BPTF-dependent loss of accessibility and a Set1-dependent H3K4me3 peak showed the strongest overall correlation with transcription. Finally, we observed that knockdown of *bptf* replicates many of the physiological, cellular, and molecular changes seen in *set1(RNAi)* planarians, including the inability of their stem cells to repopulate after treatment with ionizing radiation.

At the same time, our data also indicate that the local H3K4me3 density installed by each methyltransferase is not the only determinant of genomic output and that there are additional unknown features at Set1 and MLL1/2 target genes that distinguish them from other loci with “wide” and “narrow” H3K4me3 peaks. We observed interesting changes in *bptf(RNAi)* stem cells that were independent of Set1 and/or H3K4me3. For example, *bptf(RNAi)* stem cells exhibit an apparent dysregulation of chromatin and transcriptional regulation. Notably, several of the H3K4me3- independent genes regulated by BPTF/NuRF are other chromatin regulatory proteins, particularly those associated with gene silencing (**Figure 7, Supplemental Figure S5**). Although many future studies will be needed to untangle the multilayered downstream effects of these changes, these findings support the hypothesis that chromatin regulation is an essential and non-redundant part of the mechanism stem cells use to maintain their identity and function in vivo.

Our study is also important because the context dependence and precision of NuRF activity likely informs its biological roles in vivo. In humans, BPTF appears to have dose-dependent functions: haploinsufficiency of BPTF causes abnormal craniofacial and brain developmental [52, 53], whereas over-expression of BPTF is linked to oncogenic behavior in various cancers [54–56]. These data suggest that some cell types and/or genomic loci are more sensitive to the amount of BPTF-dependent NuRF activity than others. However, the exact mechanisms driving these functional outcomes in different in vivo settings remain unclear. Because we robustly reduced, but did not eliminate, *bptf* expression, we identified the loci and genes that are most sensitive to precise levels of NuRF activity in planarian stem cells. Despite this incomplete loss, *bptf(RNAi)* animals have severe and penetrant phenotypes, showing that normal NuRF activity levels are critical for in vivo cellular function in planarians as well.

## Limitations of the study

When interpreting our data, several points must be considered. First, we acknowledge that the method for isolating stem cells from planarians is based on DNA content, *i.e.*, cells with >2n DNA [41]. This means that most cells in our study, and most molecular/genomic studies of planarian stem cells, are in the late S/G2/M phases. It is possible that this characteristic may skew our chromatin accessibility data (ATAC-seq) toward a less accessible state, as chromosomes become highly compacted in M phase and studies using synchronized cells have reported decreased ATAC-seq signal as cells progress through these phases of cell cycle [57]. In this study, we are comparing cells in which we have depleted an integral subunit of a chromatin remodeling complex, BPTF, to control cells. Despite the possible dampening of ATAC-seq signal due to cell cycle compaction, we still detect many significant changes in chromatin accessibility. However, it is possible that we would detect many more if we could isolate all stem cells, including those with 2n DNA content (in G1 or G0).

Second, knockdown of *bptf* may cause a small, but consistent, loss of accessibility at many gene promoters, but these changes may not reach the threshold required by csaw to identify them as differentially accessible (DA). Related, some gene loci may have highly dynamic chromatin states, making them difficult to assay their accessibility reproducibly across replicates. Both scenarios could depress our ability to detect significant changes in chromatin accessibility in *bptf(RNAi)* stem cells, which then also affects our correlation analyses with gene expression. This issue is not unique to planarian studies, as similarly weak correlations between ATAC-seq and RNA-seq changes have been reported in other species and contexts [58]. Yet it reinforces the importance of considering the limitations of these assays.

Finally, the planarian field is still in the early stages of understanding the genome(s) of these animals and how they are regulated. Only recently were two chromosome-level assemblies of the *S.med* genome released [43, 59], and work remains to be done to understand its regulatory networks. For example, although the *S.med* genome is relatively large (∼800MB) with sizeable regions of repetitive sequence, the genic regions are often very dense. These details are relevant to any genomic assays, as they impact steps such as mapping ATAC-seq peaks to their nearest gene model. However, this is an issue for all genomes and genomic analyses to different extents. Nevertheless, we anticipate that our data will provide a meaningful contribution to the characterization of genome loci that regulate planarian stem cell function.

## METHODS

### Homology predictions and analyses

SmedBPTF homologs were identified by tBLASTn searches against the planarian genome [43] with the protein sequences of the human BPTF (NP_872579.2) and fly NURF301 (Q9W0T1.2) proteins, then confirmed by reciprocal BLAST to the SwissProt database. The SMART sequence analysis tool [60, 61] was used to identify conserved domains in all three BPTF/NURF301 homologs (http://smart.embl-heidelberg.de/).

### Gene cloning and RNAi

Two separate, non-overlapping regions of the identified Smed-bptf gene were cloned from a cDNA library using the following gene-specific primers: bptf-1_F= TGAGTAATGTCAATATGAAACC, bptf-1_R= AATAGTCCACATCCGTATATCT, bptf-2_F= AATGTGATAAAACACACGATAC, bptf-2_R=GAAGTTCAGTTAAAAAGTAGGC. Each PCR product was cloned into the pR-T4P vector as previously described [62]. dsRNA-containing food for RNAi treatment was generated as previously described [63]; briefly, cloned genes were transformed into bacterial strain Ht115, induced to express dsRNA with 1mM (final) IPTG, incubated with shaking at 37C for 2 hours, then pelleted, rinsed, and mixed with calf liver paste for ingestion. Each non-overlapping bptf-RNAi construct (1 & 2) induced the same phenotype in multiple experiments, after which the bptf-1 RNAi construct was used for all genomic/transcriptomic experiments.

### Planarian culture and Radiation

An asexual strain (CIW4) of *Schmidtea mediterranea* worms were maintained as described [64] with gentamicin supplemented at 50ug/ml. Worms used in experiments were starved ≥7 days prior to selection for experiments. Irradiation was done with a Gammacell 40 Exactor (MDS Nordion) at a dose rate of 85 Rad/minute.

### Live worm imaging and Whole-mount in situ hybridization (WISH)

Live worms images were captured using a Leica M205-FA. Riboprobe for *piwi-1* [65] was made by in vitro transcription with T7 polymerase from the pPR-T4P vector as previously described using DIG labeling [66]. Whole-mount in situ hybridization (WISH), including tyramide amplification of the fluorescent *piwi-1* riboprobe, was performed as previously described [67]. Worms were then mounted in modified ScaleA2 (30% glycerol/4M urea/0.1% TritonX-100/2.5% DABCO) and imaged using a confocal microscope (LSM 700, Zeiss). Quantitation of *piwi-1*^+^ cells in the confocal images was performed using ImageJ [68].

### Flow cytometry

Animals were dissociated in cold calcium- and magnesium-free buffer with 1% BSA (CMFB), filtered to remove non-dissociated tissue, pelleted, resuspended in room temperature CMFB + Hoechst 33342 (10 μg/ml), and incubated at room temperature in the dark for 1.5 hours. Propidium iodide was added (5 μg/ml) for the last 5 minutes to assess cell viability. Viable >2n “X1” stem cells [41] were then collected using a MoFlo Legacy (Beckman Coulter) or BD Influx (BD Biosciences).

### ATAC-seq analysis

Two replicates of 50K stem cells per *control(RNAi)* and *bptf(RNAi)* worms were isolated by flow cytometry and subjected to ATAC as previously described [42]. The resulting ATAC-seq libraries were sequenced on the Illumina NextSeq 500 (paired-end reads) and quality check (QC) was done using fastqc (version 0.11.9) [69]. Fragment size distributions were generated using ATACseqQC (version 1.26.0) with the fragSizeDist function [70]. Bowtie2 (version 2.4.4) [71] was used to align sample reads to the S3h1 *Schmidtea mediterranea* genome [43] using the parameters --very-sensitive -X 1000. After alignment, samtools [72] was used to create BAM files for each sample with those alignments that are properly paired (samtools view -h -b -f 3). Samples were then subsampled based on library complexity and processed as previously described [73]. PCR duplicated reads were removed using the MarkDuplicates function in Picard (version 2.26.4) (http://broadinstitute.github.io/picard). To compensate for Tn5 insertion, a Tn5 shift (+4 and -5 bp) was applied using the bedpeTn5shift.sh script [73]. ATAC-seq peaks were then called using the MACS2 [74] callpeak function (version 2.2.7.1) and parameters -g 8.40e8 -q 0.05 --keep-dup all. Consensus peaks present in both replicates were identified using BEDtools [75] and the annotation of MACS2 peaks to gene models was done using ChIPSeeker (version 1.38.0) [76, 77]. deepTools (version 3.5.1) [78] was used to generate normalized coverage files (bigWig files) and profile plots, using parameters bamCoverage --binSize 5 --smoothLength 60 -- normalizeUsing RPKM. The Integrative Genomics Viewer (IGV version 2.16.2) was used to visualize bigWig files [79]. The identification of differentially accessible ATAC-seq peaks was done using csaw (version 1.36.0) [80] as recommended in a previous study [73]. The GO (gene ontology) enrichment analysis was done using enricher function from package clusterProfiler (version 4.10.0) (14, 15). Further quantitative and statistical analysis was performed using R (version 4.3.1) and figures were created using ggplot2 [81].

### Gene expression analysis

Four replicates of 100K X1 stem cells per *control(RNAi)* and *bptf(RNAi)* condition per time point were isolated by flow cytometry (see above) and collected directly into Trizol regent. The cells were then homogenized and total RNA extracted per the manufacturer protocol. RNAseq libraries were then generated using the Illumina TruSeq kit and sequenced in 50bp single reads using the Illumina HiSeq 2500. Single-end reads were then aligned to the S3h1 *Schmidtea mediterranea* genome using Hisat2 (version 2.2.1) [82, 83] and transcript quantification was performed using StringTie (version 2.1.7). Differential gene expression analysis was then accomplished using DESeq2 (version 1.42.0) [84, 85]. Genes were considered differentially expressed if they had p-adj < 0.05. Further quantitative and statistical analysis was performed using R (version 4.3.1) and figures were created using ggplot2 [81].

### Chromatin-immunoprecipitation plus DNA sequencing (ChIP-seq)

Chromatin-immunoprecipitation was performed as previously described [22, 86].For each replicate/condition, 300K planarian X1 stem cells were isolated by FACS (see above), cross-linked in 4% PFA, and mixed with 10M (cross-linked) Drosophila S2 cells before proceeding to the next steps. Chromatin was sheared in the S220 Focused Ultrasonicator (Covaris) with silia beads and immunoprecipitation was performed with an antibody to H3K4me3 (Cell Signaling #9751). Isolated genomic DNA from both input and ChIP samples were used to generate libraries using the Illumina TruSeq kit and then sequenced (single end) on the Illumina HiSeq 2500. Quality checks of all resulting reads (QC) were done using fastqc (version 0.11.9) [69] followed by alignment to the S3h1 *Schmidtea mediterranea* genome with bowtie2 (version 2.4.4) [83, 87]. PCR duplicated reads were removed using MarkDuplicates function in Picard (version 2.26.4) (http://broadinstitute.github.io/picard). Peak calling was done using the MACS2 [74] callpeak function (version 2.2.7.1) with –g 8.40e8 --nomodel and -q 0.01 parameters. Differential analysis between *set1(RNAi), mll1/2(RNAi),* and *bptf(RNAi)* H3K4me3 ChIP libraries with their individual matching *control(RNAi)* libraries was performed using diffReps (version 1.55.6), paramaters --btr --bco --nsd --frag 250 [88]. Consensus peaks present in both replicates were identified using BEDtools [75] and peaks annotation to gene models was done using ChIPSeeker (version 1.38.0) [76, 77]. deepTools (version 3.5.1) [78] was used to generate normalized coverage files (bigWig files) and for creating profile plots (bamCoverage --normalizeUsing RPKM). Integrative Genomics Viewer (IGV version 2.16.2) was used to visualize bigWig files [79]. The GO (gene ontology) enrichment analysis was done using enricher function from package clusterProfiler (version 4.10.0) [89, 90]. Further quantitative and statistical analysis was performed using R (version 4.3.1) and figures were created using ggplot2 [81].

### GEO accession numbers

The Bioproject number for all new genomic sequencing data reported here is: **PRJNA1093065**. Reanalyzed H3K4me3 ChIP-seq data from *set1(RNAi)* and *mll1/2(RNAi)* stem cells (and their matching control stem cells) was previously published [22] and can be found under GEO accession number GSE74153.

## SUPPLEMENTAL INFORMATION

Supplemental information includes five figures of supporting data and three tables: Table S1 = differential ATAC-seq peaks identified by csaw,

Table S2= differential gene expression calculated by DEseq2, and

Table S3= Set1 and MLL1/2 target loci identified by DiffReps, based on reanalyzes of previously reported H3K4me3-ChIP data (GEO GSE74153) [22]using the S3h1 genome assembly [43].

## AUTHOR CONTRIBUTIONS

E.M.D. designed and performed all experimental work and wrote the manuscript; P.V. performed all bioinformatic analyses and contributed extensively to discussions of results and interpretation of findings. A.S.A. contributed significantly to discussions of results, interpretation of findings, and revising of the manuscript.

## ACKNOWLEDGEMENTS

The authors would like to thank the Cytometry, Molecular Biology, Imaging, and Tissue Culture cores at the Stowers Institute for Medical Research for all their hard work in maintaining such well-run facilities and in assisting E.M. Duncan as she collected and/or processed samples. We would also like to thank Laura Banaszynski, Ann Morris and Eric Ross for their valuable feedback on the manuscript. This work was funded by NIGMS grants 5R35GM142679 (E.M.D.) and 5R37GM057260 (A.S.A.).

## REFERENCES

1. Elliott SA, Alvarado AS: Planarians and the History of Animal Regeneration: Paradigm Shifts and Key Concepts in Biology. Methods Mol Biol 2018, 1774:207–239.

2. Eisenhoffer GT, Kang H, Sanchez Alvarado A: Molecular analysis of stem cells and their descendants during cell turnover and regeneration in the planarian Schmidtea mediterranea. Cell Stem Cell 2008, 3:327–339.

3. Sahu S, Dattani A, Aboobaker AA: Secrets from immortal worms: What can we learn about biological ageing from the planarian model system? Semin Cell Dev Biol 2017, 70:108–121.

4. Wagner DE, Wang IE, Reddien PW: Clonogenic neoblasts are pluripotent adult stem cells that underlie planarian regeneration. Science 2011, 332:811–816.

5. Oviedo NJ, Pearson BJ, Levin M, Sanchez Alvarado A: **Planarian PTEN homologs regulate stem cells and regeneration through TOR signaling**. Dis Model Mech 2008, 1:131–143; discussion 141.

6. Pearson BJ, Sanchez Alvarado A: A planarian p53 homolog regulates proliferation and self-renewal in adult stem cell lineages. Development 2010, 137:213–221.

7. Swapna LS, Molinaro AM, Lindsay-Mosher N, Pearson BJ, Parkinson J: Comparative transcriptomic analyses and single-cell RNA sequencing of the freshwater planarian Schmidtea mediterranea identify major cell types and pathway conservation. Genome Biol 2018, 19:124.

8. Garcia-Castro H, Kenny NJ, Iglesias M, Alvarez-Campos P, Mason V, Elek A, Schonauer A, Sleight VA, Neiro J, Aboobaker A, et al: ACME dissociation: a versatile cell fixation-dissociation method for single-cell transcriptomics. Genome Biol 2021, 22:89.

9. Zeng A, Li H, Guo L, Gao X, McKinney S, Wang Y, Yu Z, Park J, Semerad C, Ross E, et al: Prospectively Isolated Tetraspanin(+) Neoblasts Are Adult Pluripotent Stem Cells Underlying Planaria Regeneration. Cell 2018, 173:1593–1608 e1520.

10. van Wolfswinkel JC, Wagner DE, Reddien PW: Single-cell analysis reveals functionally distinct classes within the planarian stem cell compartment. Cell Stem Cell 2014, 15:326–339.

11. Benham-Pyle BW, Brewster CE, Kent AM, Mann FG, Jr., Chen S, Scott AR, Box AC, Sanchez Alvarado A: Identification of rare, transient post-mitotic cell states that are induced by injury and required for whole-body regeneration in Schmidtea mediterranea. Nat Cell Biol 2021, 23:939–952.

12. Fincher CT, Wurtzel O, de Hoog T, Kravarik KM, Reddien PW: **Cell type transcriptome atlas for the planarian Schmidtea mediterranea**. Science 2018, 360.

13. Plass M, Solana J, Wolf FA, Ayoub S, Misios A, Glazar P, Obermayer B, Theis FJ, Kocks C, Rajewsky N: **Cell type atlas and lineage tree of a whole complex animal by single-cell transcriptomics**. Science 2018, 360.

14. Adler CE, Seidel CW, McKinney SA, Sanchez Alvarado A: Selective amputation of the pharynx identifies a FoxA-dependent regeneration program in planaria. Elife 2014, 3:e02238.

15. Cowles MW, Omuro KC, Stanley BN, Quintanilla CG, Zayas RM: COE loss-of-function analysis reveals a genetic program underlying maintenance and regeneration of the nervous system in planarians. PLoS Genet 2014, 10:e1004746.

16. Tu KC, Cheng LC, H TKV, Lange JJ, McKinney SA, Seidel CW, Sanchez Alvarado A: **Egr-5 is a post-mitotic regulator of planarian epidermal differentiation**. Elife 2015, 4:e10501.

17. Soshnev AA, Josefowicz SZ, Allis CD: Greater Than the Sum of Parts: Complexity of the Dynamic Epigenome. Mol Cell 2016, 62:681–694.

18. Taverna SD, Li H, Ruthenburg AJ, Allis CD, Patel DJ: How chromatin-binding modules interpret histone modifications: lessons from professional pocket pickers. Nat Struct Mol Biol 2007, 14:1025–1040.

19. Wang Y, Wysocka J, Perlin JR, Leonelli L, Allis CD, Coonrod SA: **Linking covalent histone modifications to epigenetics: the rigidity and plasticity of the marks**. Cold Spring Harb Symp Quant Biol 2004, 69:161–169.

20. Barski A, Cuddapah S, Cui K, Roh TY, Schones DE, Wang Z, Wei G, Chepelev I, Zhao K: **High-resolution profiling of histone methylations in the human genome**. Cell 2007, 129:823–837.

21. Strahl BD, Ohba R, Cook RG, Allis CD: Methylation of histone H3 at lysine 4 is highly conserved and correlates with transcriptionally active nuclei in Tetrahymena. Proc Natl Acad Sci U S A 1999, 96:14967–14972.

22. Duncan EM, Chitsazan AD, Seidel CW, Sanchez Alvarado A: Set1 and MLL1/2 Target Distinct Sets of Functionally Different Genomic Loci In Vivo. Cell Rep 2015, 13:2741–2755.

23. Ruthenburg AJ, Allis CD, Wysocka J: Methylation of lysine 4 on histone H3: intricacy of writing and reading a single epigenetic mark. Mol Cell 2007, 25:15–30.

24. Wysocka J, Swigut T, Xiao H, Milne TA, Kwon SY, Landry J, Kauer M, Tackett AJ, Chait BT, Badenhorst P, et al: A PHD finger of NURF couples histone H3 lysine 4 trimethylation with chromatin remodelling. Nature 2006, 442:86–90.

25. Xiao H, Sandaltzopoulos R, Wang HM, Hamiche A, Ranallo R, Lee KM, Fu D, Wu C: Dual functions of largest NURF subunit NURF301 in nucleosome sliding and transcription factor interactions. Mol Cell 2001, 8:531–543.

26. Hamiche A, Sandaltzopoulos R, Gdula DA, Wu C: ATP-dependent histone octamer sliding mediated by the chromatin remodeling complex NURF. Cell 1999, 97:833–842.

27. Tsukiyama T, Wu C: Purification and properties of an ATP-dependent nucleosome remodeling factor. Cell 1995, 83:1011–1020.

28. Howe FS, Fischl H, Murray SC, Mellor J: **Is H3K4me3 instructive for transcription activation?** Bioessays 2017, 39:1–12.

29. Policarpi C, Dabin J, Hackett JA: Epigenetic editing: Dissecting chromatin function in context. Bioessays 2021, 43:e2000316.

30. Tanny JC, Erdjument-Bromage H, Tempst P, Allis CD: Ubiquitylation of histone H2B controls RNA polymerase II transcription elongation independently of histone H3 methylation. Genes Dev 2007, 21:835–847.

31. Denissov S, Hofemeister H, Marks H, Kranz A, Ciotta G, Singh S, Anastassiadis K, Stunnenberg HG, Stewart AF: **Mll2 is required for H3K4 trimethylation on bivalent promoters in embryonic stem cells, whereas Mll1 is redundant**. Development 2014, 141:526–537.

32. Bledau AS, Schmidt K, Neumann K, Hill U, Ciotta G, Gupta A, Torres DC, Fu J, Kranz A, Stewart AF, Anastassiadis K: The H3K4 methyltransferase Setd1a is first required at the epiblast stage, whereas Setd1b becomes essential after gastrulation. Development 2014, 141:1022–1035.

33. Glaser S, Schaft J, Lubitz S, Vintersten K, van der Hoeven F, Tufteland KR, Aasland R, Anastassiadis K, Ang SL, Stewart AF: **Multiple epigenetic maintenance factors implicated by the loss of Mll2 in mouse development**. Development 2006, 133:1423–1432.

34. Yu BD, Hess JL, Horning SE, Brown GA, Korsmeyer SJ: **Altered Hox expression and segmental identity in Mll-mutant mice**. Nature 1995, 378:505–508.

35. Hallson G, Hollebakken RE, Li T, Syrzycka M, Kim I, Cotsworth S, Fitzpatrick KA, Sinclair DA, Honda BM: **dSet1 is the main H3K4 di- and tri-methyltransferase throughout Drosophila development**. Genetics 2012, 190:91–100.

36. **Breen TR**, **Harte PJ:** Trithorax regulates multiple homeotic genes in the bithorax and Antennapedia complexes and exerts different tissue-specific, parasegment-specific and promoter-specific effects on each. **Development** 1993, 117:119-134.

37. Hubert A, Henderson JM, Ross KG, Cowles MW, Torres J, Zayas RM: Epigenetic regulation of planarian stem cells by the SET1/MLL family of histone methyltransferases. Epigenetics 2013, 8:79–91.

38. Camacho C, Coulouris G, Avagyan V, Ma N, Papadopoulos J, Bealer K, Madden TL: **BLAST+: architecture and applications**. BMC Bioinformatics 2009, 10:421.

39. Li H, Fischle W, Wang W, Duncan EM, Liang L, Murakami-Ishibe S, Allis CD, Patel DJ: Structural basis for lower lysine methylation state-specific readout by MBT repeats of L3MBTL1 and an engineered PHD finger. Mol Cell 2007, 28:677–691.

40. Reddien PW, Bermange AL, Murfitt KJ, Jennings JR, Sanchez Alvarado A: Identification of genes needed for regeneration, stem cell function, and tissue homeostasis by systematic gene perturbation in planaria. Dev Cell 2005, 8:635–649.

41. Hayashi T, Asami M, Higuchi S, Shibata N, Agata K: Isolation of planarian X-ray-sensitive stem cells by fluorescence-activated cell sorting. Dev Growth Differ 2006, 48:371–380.

42. Buenrostro JD, Wu B, Chang HY, Greenleaf WJ: ATAC-seq: A Method for Assaying Chromatin Accessibility Genome-Wide. Curr Protoc Mol Biol 2015, 109:21 29 21-21 29 29.

43. Ivankovic M, Brand, J.N., Pandolfini, L., Brown, T., Pippel, M., Rozanski, A., Schubert, T., Grohme, M.A., Winkler, S., Robledillo, L., Zhang, M., Codino, A., Gustincich, S., Vila-Farré, M., Zhang, S., Papantonis, A., Marques, A., Rink, J. C.: A comparative analysis of planarian genomes reveals regulatory conservation in the face of rapid structural divergence. bioRxiv 2023.

44. Lun AT, Smyth GK: csaw: a Bioconductor package for differential binding analysis of ChIP-seq data using sliding windows. Nucleic Acids Res 2016, 44:e45.

45. Kharchenko PV, Alekseyenko AA, Schwartz YB, Minoda A, Riddle NC, Ernst J, Sabo PJ, Larschan E, Gorchakov AA, Gu T, et al: Comprehensive analysis of the chromatin landscape in Drosophila melanogaster. Nature 2011, 471:480–485.

46. Li JJ, Huang H, Bickel PJ, Brenner SE: Comparison of D. melanogaster and C. elegans developmental stages, tissues, and cells by modENCODE RNA-seq data. Genome Res 2014, 24:1086–1101.

47. Duncan EM, Allis CD: Errors in erasure: links between histone lysine methylation removal and disease. Prog Drug Res 2011, 67:69–90.

48. Verma P, Waterbury CKM, Duncan EM: Set1 Targets Genes with Essential Identity and Tumor-Suppressing Functions in Planarian Stem Cells. Genes (Basel) 2021, 12.

49. Badenhorst P, Voas M, Rebay I, Wu C: Biological functions of the ISWI chromatin remodeling complex NURF. Genes Dev 2002, 16:3186–3198.

50. Chen K, Chen Z, Wu D, Zhang L, Lin X, Su J, Rodriguez B, Xi Y, Xia Z, Chen X, et al: Broad H3K4me3 is associated with increased transcription elongation and enhancer activity at tumor-suppressor genes. Nat Genet 2015, 47:1149–1157.

51. Benayoun BA, Pollina EA, Ucar D, Mahmoudi S, Karra K, Wong ED, Devarajan K, Daugherty AC, Kundaje AB, Mancini E, et al: H3K4me3 Breadth Is Linked to Cell Identity and Transcriptional Consistency. Cell 2015, 163:1281–1286.

52. Stankiewicz P, Khan TN, Szafranski P, Slattery L, Streff H, Vetrini F, Bernstein JA, Brown CW, Rosenfeld JA, Rednam S, et al: Haploinsufficiency of the Chromatin Remodeler BPTF Causes Syndromic Developmental and Speech Delay, Postnatal Microcephaly, and Dysmorphic Features. Am J Hum Genet 2017, 101:503–515.

53. Glinton KE, Hurst ACE, Bowling KM, Cristian I, Haynes D, Adstamongkonkul D, Schnappauf O, Beck DB, Brewer C, Parikh AS, et al: Phenotypic expansion of the BPTF-related neurodevelopmental disorder with dysmorphic facies and distal limb anomalies. Am J Med Genet A 2021, 185:1366–1378.

54. Gong YC, Liu DC, Li XP, Dai SP: **BPTF biomarker correlates with poor survival in human NSCLC**. Eur Rev Med Pharmacol Sci 2017, 21:102–107.

55. Pan Y, Yuan F, Li Y, Wang G, Lin Z, Chen L: Bromodomain PHD-finger transcription factor promotes glioma progression and indicates poor prognosis. Oncol Rep 2019, 41:246–256.

56. Miao J, Zhang M, Huang X, Xu L, Tang R, Wang H, Han S: Upregulation of bromodomain PHD finger transcription factor in ovarian cancer and its critical role for cancer cell proliferation and survival. Biochem Cell Biol 2021, 99:304–312.

57. Yu Q, Liu X, Fang J, Wu H, Guo C, Zhang W, Liu N, Jiang C, Sha Q, Yuan X, et al: Dynamics and regulation of mitotic chromatin accessibility bookmarking at single-cell resolution. Sci Adv 2023, 9:eadd2175.

58. Kiani K, Sanford EM, Goyal Y, Raj A: Changes in chromatin accessibility are not concordant with transcriptional changes for single-factor perturbations. Mol Syst Biol 2022, 18:e10979.

59. Guo L, Bloom JS, Dols-Serrate D, Boocock J, Ben-David E, Schubert OT, Kozuma K, Ho K, Warda E, Chui C, et al: Island-specific evolution of a sex-primed autosome in a sexual planarian. Nature 2022, 606:329–334.

60. Schultz J, Milpetz F, Bork P, Ponting CP: SMART, a simple modular architecture research tool: identification of signaling domains. Proc Natl Acad Sci U S A 1998, 95:5857–5864.

61. Letunic I, Doerks T, Bork P: **SMART: recent updates, new developments and status in** 2015. Nucleic Acids Res 2015, 43:D257–260.

62. Adler CE, Alvarado AS: Systemic RNA Interference in Planarians by Feeding of dsRNA Containing Bacteria. Methods Mol Biol 2018, 1774:445–454.

63. Gurley KA, Rink JC, Sánchez Alvarado A: β-Catenin Defines Head Versus Tail Identity During Planarian Regeneration and Homeostasis. Science 2008, 319:323–327.

64. Newmark PA, Sánchez Alvarado A: Bromodeoxyuridine Specifically Labels the Regenerative Stem Cells of Planarians. Developmental Biology 2000, 220:142–153.

65. Reddien PW, Oviedo NJ, Jennings JR, Jenkin JC, Sanchez Alvarado A: **SMEDWI-2 is a PIWI-like protein that regulates planarian stem cells**. Science 2005, 310:1327–1330.

66. Pearson BJ, Eisenhoffer GT, Gurley KA, Rink JC, Miller DE, Sánchez Alvarado A: **Formaldehyde-based whole-mount in situ hybridization method for planarians**. Developmental dynamics : an official publication of the American Association of Anatomists 2009, 238:443–450.

67. King RS, Newmark PA: In situ hybridization protocol for enhanced detection of gene expression in the planarian Schmidtea mediterranea. BMC Developmental Biology 2013, 13:8.

68. Schindelin J, Arganda-Carreras I, Frise E, Kaynig V, Longair M, Pietzsch T, Preibisch S, Rueden C, Saalfeld S, Schmid B, et al: Fiji: an open-source platform for biological-image analysis. Nat Methods 2012, 9:676-682.

69. FastQC: a quality control tool for high throughput sequence data. Available online at: [https://www.bioinformatics.babraham.ac.uk/projects/fastqc/]

70. Ou JH, Liu HB, Yu J, Kelliher MA, Castilla LH, Lawson ND, Zhu LJ: ATACseqQC: a Bioconductor package for post-alignment quality assessment of ATAC-seq data. Bmc Genomics 2018, 19.

71. Langdon WB: Performance of genetic programming optimised Bowtie2 on genome comparison and analytic testing (GCAT) benchmarks. BioData Min 2015, 8:1.

72. Li H, Handsaker B, Wysoker A, Fennell T, Ruan J, Homer N, Marth G, Abecasis G, Durbin R, Genome Project Data Processing S: **The Sequence Alignment/Map format and SAMtools**. Bioinformatics 2009, 25:2078–2079.

73. Reske JJ, Wilson MR, Chandler RL: ATAC-seq normalization method can significantly affect differential accessibility analysis and interpretation. Epigenetics Chromatin 2020, 13:22.

74. Zhang Y, Liu T, Meyer CA, Eeckhoute J, Johnson DS, Bernstein BE, Nusbaum C, Myers RM, Brown M, Li W, Liu XS: Model-based analysis of ChIP-Seq (MACS). Genome Biol 2008, 9:R137.

75. Quinlan AR: BEDTools: The Swiss-Army Tool for Genome Feature Analysis. Curr Protoc Bioinformatics 2014, 47:11 12 11-34.

76. Wang Q, Li M, Wu T, Zhan L, Li L, Chen M, Xie W, Xie Z, Hu E, Xu S, Yu G: Exploring Epigenomic Datasets by ChIPseeker. Curr Protoc 2022, 2:e585.

77. Yu G, Wang LG, He QY: ChIPseeker: an R/Bioconductor package for ChIP peak annotation, comparison and visualization. Bioinformatics 2015, 31:2382–2383.

78. Ramirez F, Dundar F, Diehl S, Gruning BA, Manke T: **deepTools: a flexible platform for exploring deep-sequencing data**. Nucleic Acids Res 2014, 42:W187–191.

79. Robinson JT, Thorvaldsdottir H, Winckler W, Guttman M, Lander ES, Getz G, Mesirov JP:**Integrative genomics viewer**. Nat Biotechnol 2011, 29:24–26.

80. Lun ATL, Smyth GK: csaw: a Bioconductor package for differential binding analysis of ChIP-seq data using sliding windows. Nucleic Acids Research 2016, 44.

81. Wickham H: ggplot2: Elegant Graphics for Data Analysis. Ggplot2: Elegant Graphics for Data Analysis 2009:1-212.

82. Kim D, Langmead B, Salzberg SL: HISAT: a fast spliced aligner with low memory requirements. Nat Methods 2015, 12:357–360.

83. Ivankovic M, Brand JN, Pandolfini L, Brown T, Pippel M, Rozanski A, Schubert T, Grohme MA, Winkler S, Robledillo L, et al: A comparative analysis of planarian genomes reveals regulatory conservation in the face of rapid structural divergence. bioRxiv 2023:2023.2012.2022.572568.

84. Love MI, Huber W, Anders S: Moderated estimation of fold change and dispersion for RNA-seq data with DESeq2. Genome Biology 2014, 15.

85. Pertea M, Pertea GM, Antonescu CM, Chang TC, Mendell JT, Salzberg SL: **StringTie enables improved reconstruction of a transcriptome from RNA-seq reads**. Nature Biotechnology 2015, 33:290-+.

86. Lee TI, Johnstone SE, Young RA: Chromatin immunoprecipitation and microarray-based analysis of protein location. Nature Protocols 2006, 1:729–748.

87. Langmead B, Salzberg SL: **Fast gapped-read alignment with Bowtie 2**. Nature Methods 2012, 9:357–U354.

88. Shen L, Shao NY, Liu X, Maze I, Feng J, Nestler EJ: diffReps: detecting differential chromatin modification sites from ChIP-seq data with biological replicates. PLoS One 2013, 8:e65598.

89. Wu TZ, Hu EQ, Xu SB, Chen MJ, Guo PF, Dai ZH, Feng TZ, Zhou L, Tang WL, Zhan L, et al: clusterProfiler 4.0: A universal enrichment tool for interpreting omics data. Innovation 2021, 2.

90. Yu GC, Wang LG, Han YY, He QY: **clusterProfiler: an R Package for Comparing Biological Themes Among Gene Clusters**. Omics-a Journal of Integrative Biology 2012, 16:284–287.

91. Li H, Ilin S, Wang W, Duncan EM, Wysocka J, Allis CD, Patel DJ: Molecular basis for site-specific read-out of histone H3K4me3 by the BPTF PHD finger of NURF. Nature 2006, 442:91–95.

92. Lei K, Thi-Kim Vu H, Mohan RD, McKinney SA, Seidel CW, Alexander R, Gotting K, Workman JL, Sanchez Alvarado A: **Egf Signaling Directs Neoblast Repopulation by Regulating Asymmetric Cell Division in Planarians**. Dev Cell 2016, 38:413–429.

