## Supplemental figures for "Chromatin remodeling protein BPTF regulates transcriptional stability in planarian stem cells"

**Verma et al, SUPPLEMENTAL FIGURES**

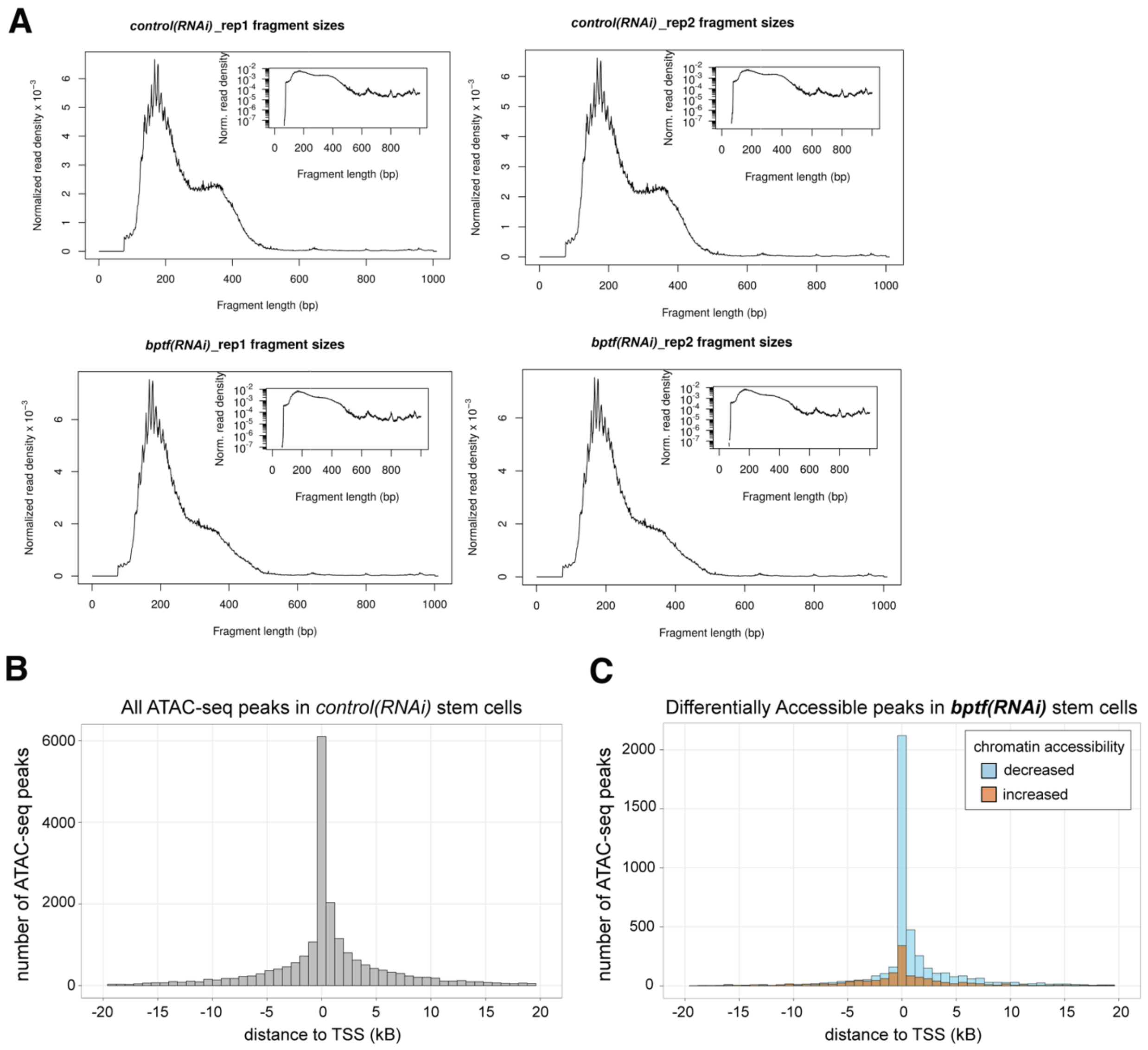


**Supplemental Figure S1. ATAC-seq uncovers significant changes in chromatin accessibility in *bptf(RNAi)* stem cells.**

A) Individual fragment size distribution plots for each ATAC-seq library; two replicates are from control(RNAi) stem cells and two from bptf(RNAi).

B) Histogram summarizing the distance to nearest TSS (transcription start site) for all MAC2-called peaks in *control(RNAi)* stem cells; 800bp binwidth.

A) Histogram summarizing the distance to TSS of all differentially accessible (DA) ATAC-seq peaks (csaw) in *bptf(RNAi)* stem cells compared to *control(RNAi)* stem cells (800bp binwidth). Blue bars = peaks with decreased accessibility (4652 peaks), orange bars = peaks with increased accessibility (1411 peaks).

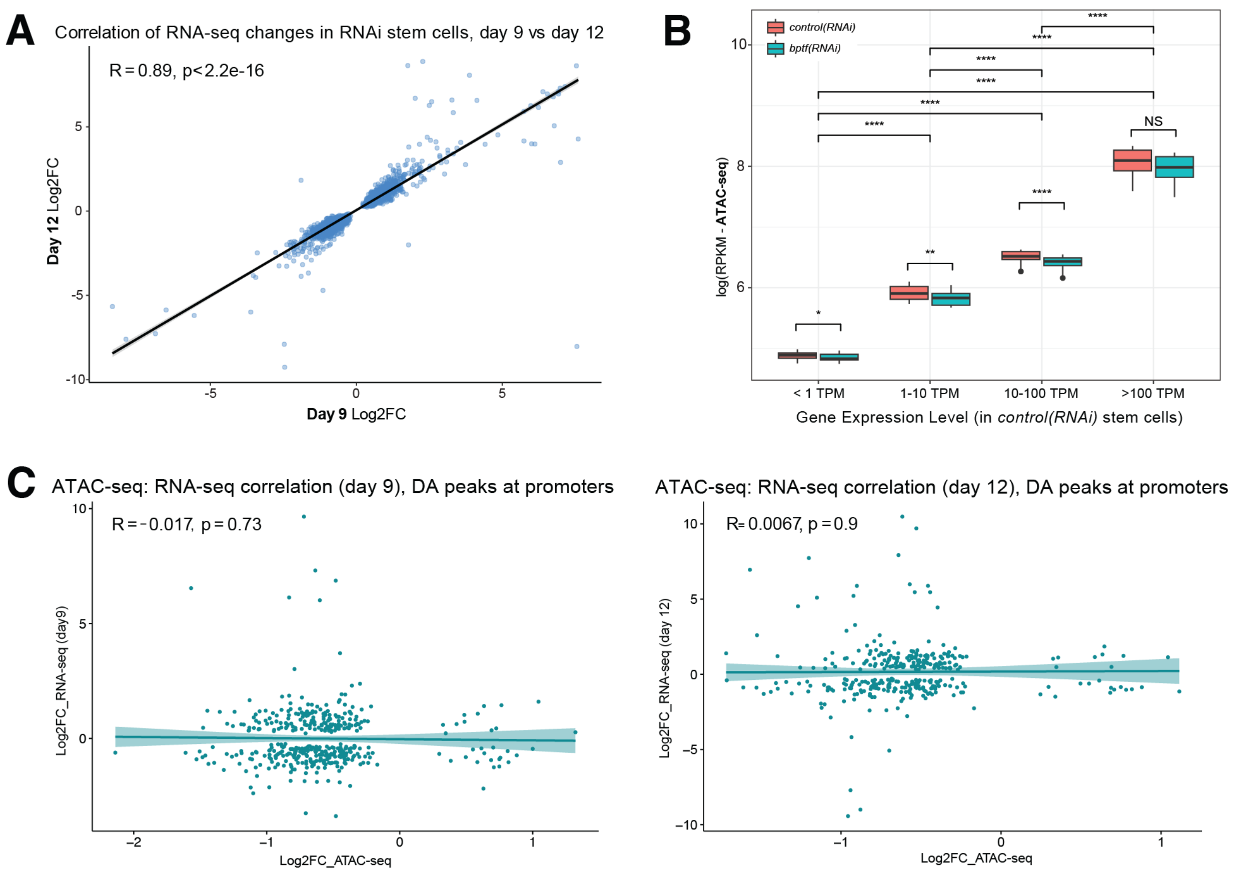


**Supplemental Figure S2. Knockdown of *bptf* leads to stable changes in gene expression in planarian stem cells.**

A) Correlation plot comparing differential gene expression (DEseq2, *bptf(RNAi)* vs *control(RNAi)*) at two time points after RNAi (day 9 and day 12)

B) Box plot of average ATAC-seq signal at genes with very low expression (<1 TPM, low expression (1-10 TPM), moderate expression (10-100 TPM), and high expression (> 100 TPM) in control stem cells. Overall, average ATAC-seq signal correlates with average expression. *Bptf(RNAi)* stem cells have lower average ATAC-seq for all groups except the highest expression bin (> 100 TPM). Statistical significance among the TPM groups was determined by the TukeyHSD test; * = p≤ 0.05, ** = p ≤ 0.01, *** = p≤ 0.001, **** = p≤ 0.0001. The Wilcoxon test was used to determine the significance between *control(RNAi) and bptf(RNAi)* signal (p-value with Bonferroni correction; * = p≤ 0.05, ** = p≤ 0.01, *** = p≤ 0.001, **** = p≤ 0.0001).

C) Correlation plots comparing log2FC changes in chromatin accessibility (ATAC-seq) for DA peaks (csaw, *bptf(RNAi)* vs *control(RNAi)*) at gene promoters with the log2FC changes (RNA-seq) for those genes (if significant). Left plot = day 9 RNA-seq, right = day 12. Statistical significance among the TPM groups was determined TukeyHSD test (* = p≤ 0.05, ** = p ≤ 0.01, *** = p≤ 0.001, **** = p≤ 0.0001) and Wilcoxon test was used to determine the significance between *control(RNAi) and bptf(RNAi)* signal (p-value with Bonferroni correction; * = p≤ 0.05, ** = p≤ 0.01, *** = p≤ 0.001, **** = p≤ 0.0001).

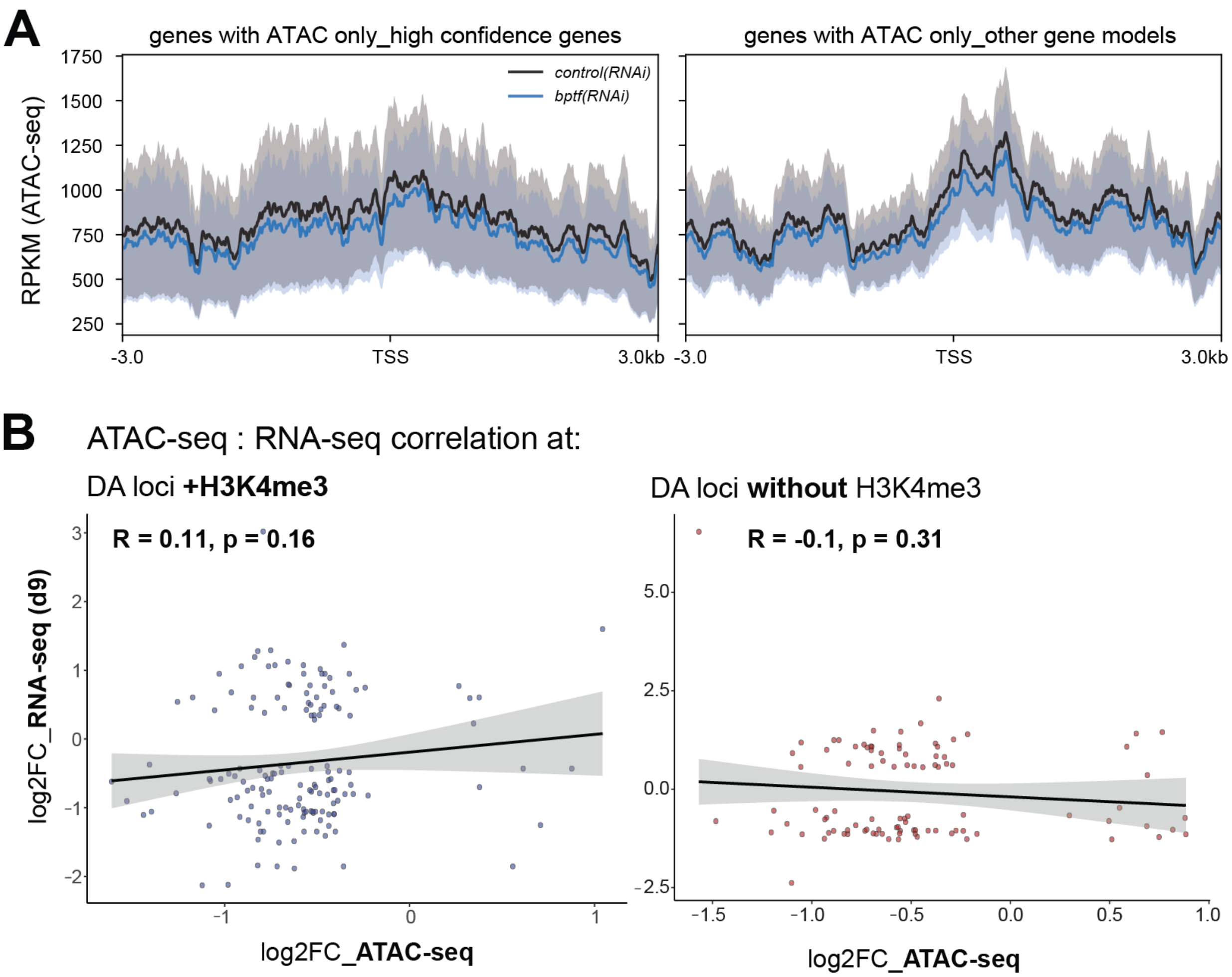


**Supplemental Figure S3. ATAC-seq signal at genes with H3K4me3 peaks is more sensitive to loss of BPTF.**

A) Profile plots of ATAC-seq signal at genes with ATAC-seq peaks only (no MACS2-called H3K4me3 peak); left plot = average across "high confidence" genes, right plot = average across rest gene models (annotations created and categorized in [1]).

B) Plots correlating changes in chromatin accessibility (log2FC ATAC-seq) with changes in transcription (log2FC RNA-seq) for those genes with an H3K4me3 peak at promoter (left plot) and those without one (right plot). RNA-seq data here is from 9 days post RNAi treatment; plots in Figure 4 are with RNA-seq data from day 12 post-RNAi.

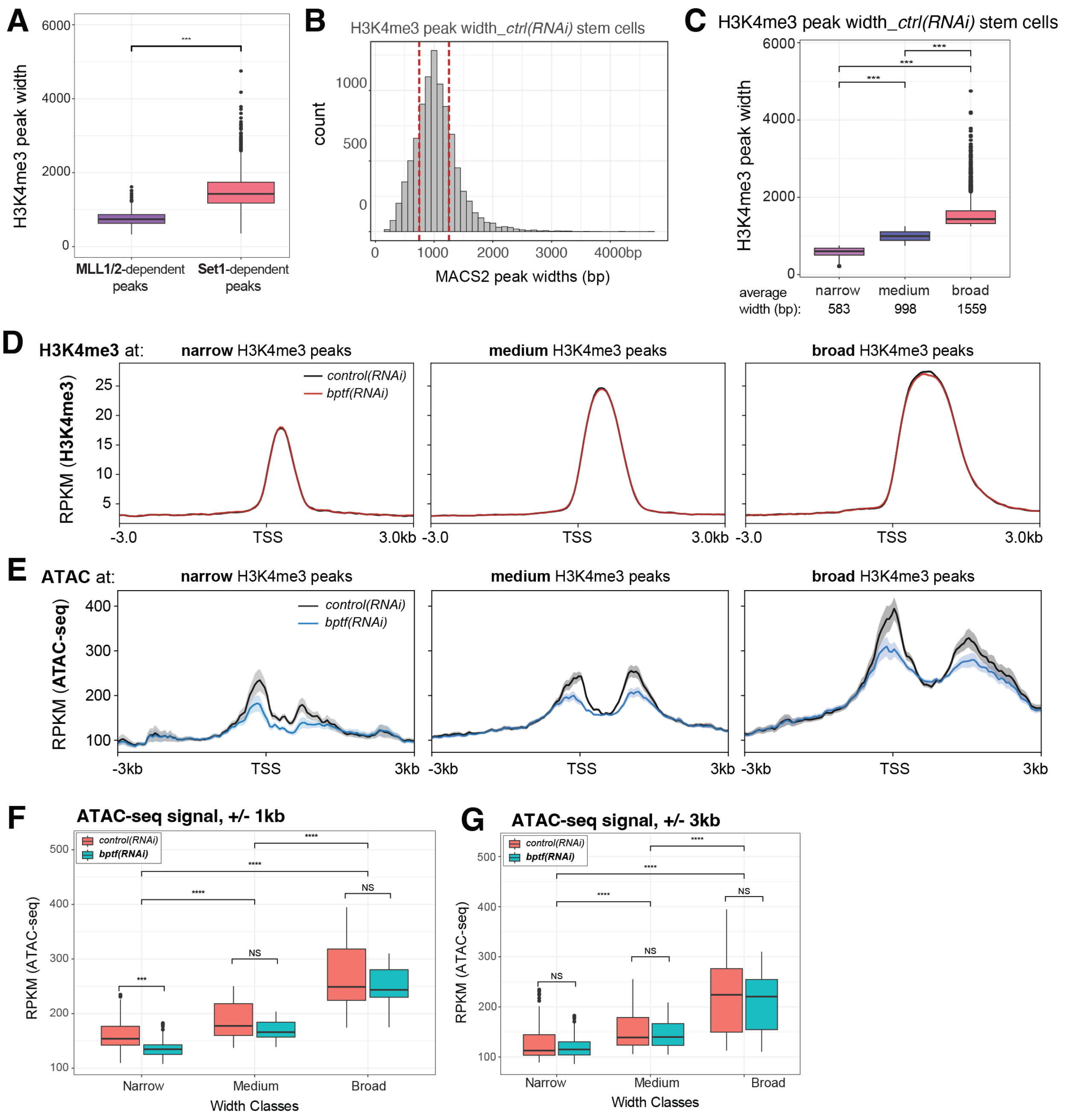


**Supplemental Figure S4. H3K4me3 peak width correlates with ATAC-seq signal.**

A) Box plot of average H3K4me3 peak width at Set1 vs MLL1/2 target genes. Statistical significance was determined using Student's t test (* = p-value ≤0.05, **= p≤ 0.01, *** = p≤ 0.001, **** = p≤ 0.0001).

B) Histogram of H3K4me3 peak width for all MACS2-called peaks in *control(RNAi)* stem cells. Dashed lines = divisions of "narrow", "medium", and "broad" groups.

C) Box plot of average H3K4me3 peak width of the peaks in each peak width group. Statistical significance was determined using TukeyHSD test (* = p≤ 0.05, ** = p≤ 0.01, *** = p≤ 0.001, **** = p≤ 0.0001)

D) Profile plots of average H3K4me3 signal at the gene loci in each peak width group.
E) Profile plots of average ATAC-seq signal at the gene loci in each peak width group.
F) Box plots quantitating the ATAC-seq signal in E, +/- 1kb. Statistical significance was determined using TukeyHSD test (* = p≤ 0.05, ** = p≤ 0.01, *** = p≤ 0.001, **** = p≤ 0.0001) and Wilcoxon test to determine the significance between *control(RNAi) and bptf(RNAi)* signal (p-value with Bonferroni correction: * = p≤ 0.05, ** = p≤ 0.01, *** = p≤ 0.001, **** p≤ 0.0001).
G) Box plots quantitating the ATAC-seq signal in E, +/- 3kb. Statistical significance was determined using TukeyHSD test (* = p≤ 0.05, ** = p≤ 0.01, *** = p≤ 0.001, **** = p≤ 0.0001) and Wilcoxon test to determine the significance between *control(RNAi) and bptf(RNAi)* signal (p-value with Bonferroni correction: * = p≤ 0.05, ** = p≤ 0.01, *** = p≤ 0.001, **** = p≤ 0.0001).

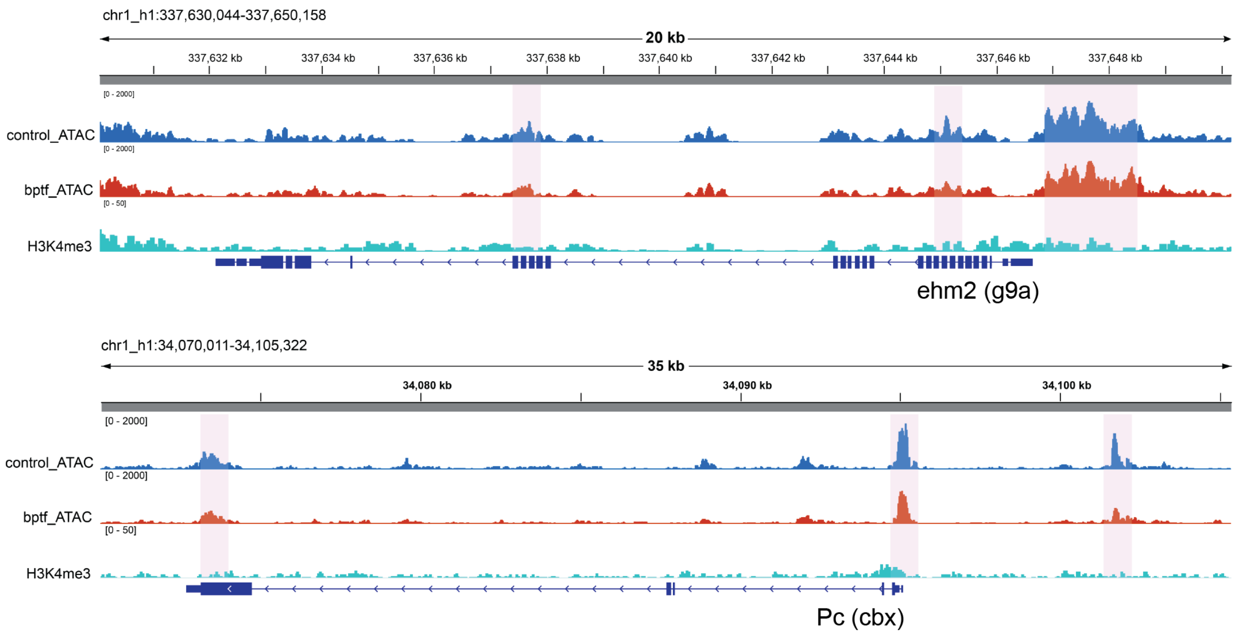


**Supplemental Figure S5. Example tracks of genes with positive correlation between ATAC-seq and RNA-seq changes.**
Representative examples of genes that have differentially accessibly peaks in *bptf(RNAi)* stem cells and positively correlating changes in RNA-seq expression (i.e., down-regulated in both), but do not have H3K4me3 peaks. Both genes shown here encode proteins associated with chromatin silencing.

**References:**

1. Ivankovic, M., Brand, J.N., Pandolfini, L., Brown, T., Pippel, M., Rozanski, A., Schubert, T., Grohme, M.A., Winkler, S., Robledillo, L., Zhang, M., Codino, A., Gustincich, S., Vila-Farré, M., Zhang, S., Papantonis, A., Marques, A., Rink, J. C., *A comparative analysis of planarian genomes reveals regulatory conservation in the face of rapid structural divergence.* bioRxiv, 2023.
